# Systematic perturbation screens decode regulators of inflammatory macrophage states and identify a role for *TNF* mRNA m6A modification

**DOI:** 10.1101/2024.04.12.589122

**Authors:** Simone M Haag, Shiqi Xie, Celine Eidenschenk, Jean-Philippe Fortin, Marinella Callow, Mike Costa, Aaron Lun, Chris Cox, Sunny Z Wu, Rachana N Pradhan, Jaclyn Lock, Julia A Kuhn, Loryn Holokai, Minh Thai, Emily Freund, Ariane Nissenbaum, Mary Keir, Christopher J Bohlen, Scott Martin, Kathryn Geiger-Schuller, Hussein A Hejase, Brian L Yaspan, Sandra Melo Carlos, Shannon J Turley, Aditya Murthy

## Abstract

Macrophages adopt dynamic cell states with distinct effector functions to maintain tissue homeostasis and respond to environmental challenges. During chronic inflammation, macrophage polarization is subverted towards sustained inflammatory states which contribute to disease, but there is limited understanding of the regulatory mechanisms underlying these disease-associated states. Here, we describe a systematic functional genomics approach that combines genome-wide phenotypic screening in primary murine macrophages with transcriptional and cytokine profiling of genetic perturbations in primary human monocyte-derived macrophages (hMDMs) to uncover regulatory circuits of inflammatory macrophage states. This process identifies regulators of five distinct inflammatory states associated with key features of macrophage function. Among these, the mRNA m6A writer components emerge as novel inhibitors of a TNFα-driven cell state associated with multiple inflammatory pathologies. Loss of m6A writer components in hMDMs enhances *TNF* transcript stability, thereby elevating macrophage TNFα production. A PheWAS on SNPs predicted to impact m6A installation on *TNF* revealed an association with cystic kidney disease, implicating an m6A-mediated regulatory mechanism in human disease. Thus, systematic phenotypic characterization of primary human macrophages describes the regulatory circuits underlying distinct inflammatory states, revealing post-transcriptional control of TNF mRNA stability as an immunosuppressive mechanism in innate immunity.

## INTRODUCTION

As key constituents of all tissue microenvironments, macrophages play essential roles in guiding tissue repair, remodeling, antimicrobial responses and adaptive immunity^1^. This versatility depends on their ability to adopt a spectrum of cellular states with distinct effector functions in response to environmental triggers^2,3^. High-resolution single cell gene expression profiling has allowed detailed mapping of myeloid cellular states, revealing macrophage subsets that are enriched in inflammatory diseases and associated with patient outcome^4–7^. However, the underlying regulatory mechanisms which establish inflammatory macrophage states are poorly characterized. Identifying these would facilitate the development of novel macrophage-centered therapies.

Single-cell CRISPR (scCRISPR) screening, a functional genomics approach which combines CRISPR-mediated gene editing with single-cell RNA-sequencing (scRNA-seq), has emerged as a powerful means to identify regulatory circuits of cell states^8,9^. While this approach has been successfully applied in primary human T cells^10,11^, its application in the study of primary human macrophages is lagging. Macrophages express abundant amounts of the central restriction factor SAMHD1 which prevents lentiviral integration^12^, and despite being readily available, human monocyte-derived macrophages (hMDMs) cannot easily be expanded, thereby limiting phenotypic screening at genome-wide scale.

Interferon-γ (IFN-γ) is one of the most potent cytokines contributing to macrophage activation. While IFN-γ induces Jak/STAT1-dependent cytokine production and antigen presentation, it also enhances macrophage responsiveness to secondary pro-inflammatory trigger such as Toll-like receptor (TLR) ligands, TNFα, or type I IFNs through a phenomenon called “priming”^13^. IFN-γ primed macrophages can adopt a spectrum of cellular states with distinct effector functions based on the nature of the secondary pro-inflammatory trigger^14^. Multiple regulatory mechanisms control these cell states, including chromatin and metabolic changes as well as paracrine and autocrine signaling networks^2,15–17^. We hypothesized that a genetic perturbation screen targeting immunosuppressive genes in IFN-γ primed macrophages would reveal negative regulators of pro-inflammatory cell states beyond a Jak/STAT1-mediated state.

Here, we describe a funneled, cross-species genomics approach combining complementary CRISPR loss-of-function screens with functional profiling of cytokine secretion in IFN-γ primed macrophages. Inflammatory macrophage states harbor distinct cytokine profiles^18,19^. Similar to the utility of cytokine profiles in defining T helper cell subsets, macrophage cytokine secretion profiles can provide an accessible and relevant readout of macrophage states, thereby facilitating the identification of the underlying regulatory circuits associated with specific functional subsets. Using this funneled approach, transcriptional profiling of human primary macrophages under genetic perturbations at single cell resolution was performed, resolving five perturbation-induced programs associated with key features of macrophage function and unique cytokine secretion profiles. Mechanistic studies pinpointed components of the mRNA m6A writer complex WTAP and ZC3H13 as potent negative regulators of a distinct TNFα cytokine-driven program in human macrophages. Specifically, loss of m6A writer components stabilized *TNF* mRNA, thus markedly elevating TNF production. Intriguingly, a PheWAS on SNPs predicted to impact m6A installation on *TNF* mRNA revealed an association with cystic kidney disease, an inflammatory pathology known to be driven by TNF^20–22^. Thus, genetic perturbation screens in primary macrophages can reveal novel immunosuppressive mechanisms relevant to human disease.

## RESULTS

### Funneled CRISPR screening of pro-inflammatory macrophage states in response to IFN-γ

To systemically elucidate regulatory circuits of pro-inflammatory macrophage states, a funneled CRISPR knockout screening approach with three tiered stages was established. First, a pooled genome-wide phenotypic screen in IFN-γ primed murine bone-marrow derived macrophages (mBMDMs), a scalable model amenable to lentiviral integration, allowed for unbiased identification of negative regulators (‘target genes’) of a proinflammatory macrophage phenotype. Next, using a non-viral gene editing strategy we recently developed^23^, an arrayed scCRISPR screen of target genes was performed in IFN-γ primed hMDMs. Finally, target genes of interest were functionally characterized in hMDMs by cytokine secretion profiling (**Fig. 1a**).

**Fig. 1:**
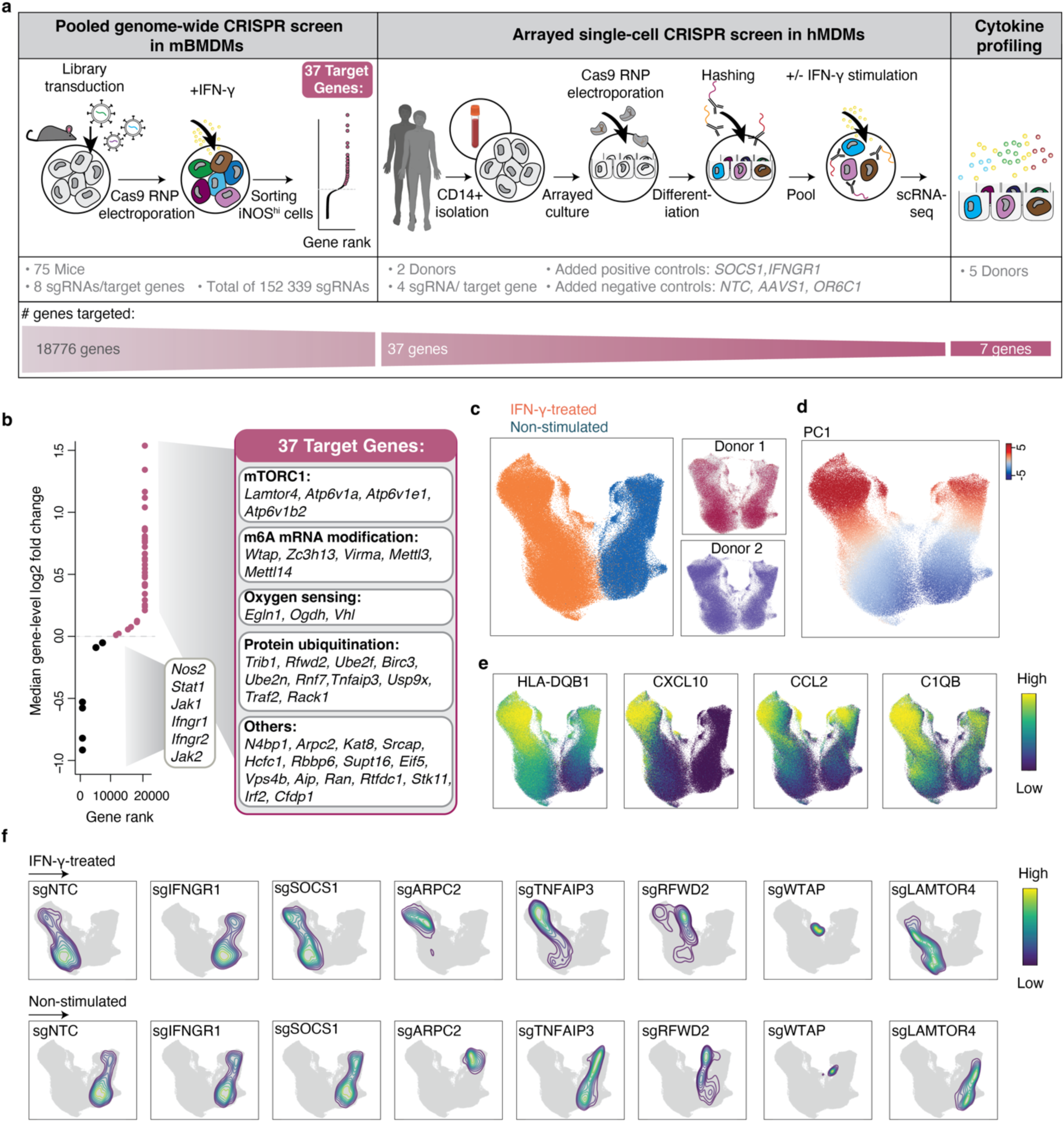
Funneled CRISPR screening approach for regulators of inflammatory macrophage states in primary cells. **a**, Study overview. Left: Genome-wide CRISPR screen in mBMDMs. Middle: Arrayed single-cell CRISPR screen in hMDMs. Right: Cytokine secretion profiling of selected target genes. **b**, Left: waterfall plot of rank-ordered normalized gene-levels of enriched sgRNAs in ‘iNOS_high’ group compared to the reference control group. Highlighted in red are 37 target genes (iNOS_high hits), FDR < 5 %. Representative sgRNAs targeting iNOS and components of the IFN-γ signaling pathway are depleted and highlighted in black. Right: 37 target genes grouped according to known biological function. **c**, Left, UMAP projection of 176,564 perturbed hMDMs post quality control and donor integration, colored by treatment. Right, UMAP projection of each of the two individual donors. **d**, PC1 in UMAP projection. **e**, UMAP with the expression of indicated genes in perturbed hMDMs. **f**, Contour density plots of hMDMs assigned to indicated sgRNA targets in UMAP space. The perturbed cell contour is shown in grayscale underneath.

To provide an appropriate phenotypic readout for the first screening stage, expression of inducible nitric oxide synthase (iNOS) was measured in IFN-γ primed mBMDMs (**Supplementary Fig. 1a)**. iNOS was chosen based on several criteria required for a robust biomarker in forward genetic screens. First, it is a stable cytosolic protein with a high dynamic range that is readily measured using fluorescence activated cell sorting (FACS). Second, priming macrophages with IFN-γ does not induce high iNOS levels; rather synergistic effects between IFN-γ (‘signal 1’) and secondary pro-inflammatory triggers (‘signal 2’) such as cytokines, innate ligands or hypoxia induce high levels of iNOS protein in mBMDMs (**Supplementary Fig. 1a, b**). Third, iNOS is broadly induced by numerous transcription factors including but not limited to NF-kB, IRF1, AP-1 and HIF1a^24,25^. Taken together, IFN-γ primed macrophages acquire diverse transcriptional states based on the nature of the synergistic trigger (‘signal 2’), which collectively promote high levels of iNOS expression (**Supplementary Fig. 1c**). Thus, iNOS was viewed as an ideal “catch all” biomarker to identify negative regulators of macrophage states downstream of multiple activation pathways.

Three independent genome-wide CRISPR loss-of-function screens were performed in mBMDMs pairing lentiviral transduction of an sgRNA library comprising 152,339 single-guide RNAs (sgRNAs) with electroporation of Cas9 protein (**Fig. 1a, left panel**). sgRNAs were isolated and sequenced from three samples: (1) library control input cells, collected three days after lentiviral transduction; (2) unstimulated reference cells, collected eleven days after lentiviral transduction and (3) IFN-γ-stimulated, iNOS high sorted cells (iNOS^hi^). The quality of all three replicates was confirmed individually and depletion of sgRNAs targeting essential genes in reference cells as compared to input cells was verified, indicating potent and consistent gene deletion efficiencies across screening replicates (**Supplementary Fig. 2a, b**). As expected, sgRNAs targeting key components of the IFN-γ signaling pathway were depleted in iNOS^hi^ cells compared to reference cells (**Fig. 1b**).

To assemble a target list of gene perturbations enriched in iNOS^hi^ cells compared to reference cells, a differential abundance analysis at the sgRNA level and two complementary gene aggregation methods across screening replicates was performed (Methods). This revealed a robust and reproducible set of 37 target genes, which grouped into 4 distinct ontologies, annotated as (1) mTORC1, (2) oxygen sensing, (3) protein ubiquitination and (4) N6-methyladenosine (m6A) mRNA modification (**Fig. 1b**). Genes not implicated in any functional category were grouped as “others”. Among the 37 genes are well-known negative regulators of inflammatory signaling such as *Tnfaip3* encoding the ubiquitin-editing enzyme A20^26^, supporting the validity of the screen. Additionally, we also identified several genes with less understood roles in immunoregulation including components of the m6A mRNA writer complex such as *Wtap* and *Zc3h13*^27,28^. Individual perturbation experiments of selected target genes confirmed increase of iNOS upon IFN-γ priming (**Supplementary Fig. 2c**).

Next, we investigated the role of these 37 genes in primary hMDMs. To overcome the resistance to viral transduction inherent to these cells, we performed scCRISPR screening in hMDMs using a recently developed non-viral gene editing strategy for efficient gene knockout^23^ coupled with single cell transcriptome readout using the 10X Genomics platform (**Fig. 1a, center panel**). CRISPR–Cas9 ribonucleoprotein (Cas9 RNPs) complexes were assembled *in vitro* and electroporated into donor blood-derived CD14+ cells in an arrayed format, with each perturbation barcoded using a combination of two antibody-conjugated hashtag oligonucleotides (HTOs). Multiplexed Cas9 RNPs of 4 sgRNAs per gene were generated for each of the 37 target genes as well as *IFNGR1* and *SOCS1*, known regulators of the IFN-γ signaling pathway, and a non-targeting control (NTC) and safe harbor genes *AAVS1* and *OR6C1,* as controls. Each perturbation was subjected to IFN-γ priming for 18 hours or left untreated. This approach generated 176,564 single cell profiles after donor integration and hashtag assignment with a median of 4,537 hMDMs per genetic perturbation (**Fig. 1c**). Uniform manifold approximation and projection (UMAP) dimensionality reduction of cellular mRNA profiles showed an even distribution of hMDMs from both donors and a distinct separation between unstimulated and IFN-γ-stimulated cells (**Fig. 1c**). A Principal Component Analysis (PCA) revealed PC1, which captured a gradient along both treatment conditions characterized by pro-inflammatory genes including components of the antigen presentation machinery (*HLA-DQB1*), inflammatory cytokines (*CCL2*, *CXCL10*) and complement system (*C1QB*) (**Figs. 1d, e and Supplementary Figs. 3a, b**). Genetic perturbations induced a shift along the PC1 gradient, indicating reconfiguration of pro-inflammatory states at basal and IFN-γ treated conditions (**Fig. 1 f)**. Scoring a monocyte and macrophage signature on the perturbed cell pool confirmed complete differentiation into mature hMDMs and excluded the possibility that the transcriptional heterogeneity was due to deficiencies in monocyte-to-macrophage differentiation (**Supplementary Fig. 3c**). While an applicable pro-inflammatory biomarker in mBMDMs, iNOS transcript levels were not detected in hMDMs likely due epigenetic silencing^29^. Given the deep transcriptional profiling of hMDMs under genetic perturbations, we next aimed to deconvolute pro-inflammatory cell states controlled by the perturbed target genes at the transcriptional level.

### Computational framework for hMDMs scCRISPR screen

To deconvolute pro-inflammatory cell states in the context of IFN-γ priming, a computational analysis in three constitutive steps was deployed: (1) exclusion of perturbations with no significant impact on gene expression; (2) identification of target gene perturbation-induced cell states and; (3) estimating regulatory circuits using transcription factor targeted genes (**Fig. 2a**).

**Fig. 2:**
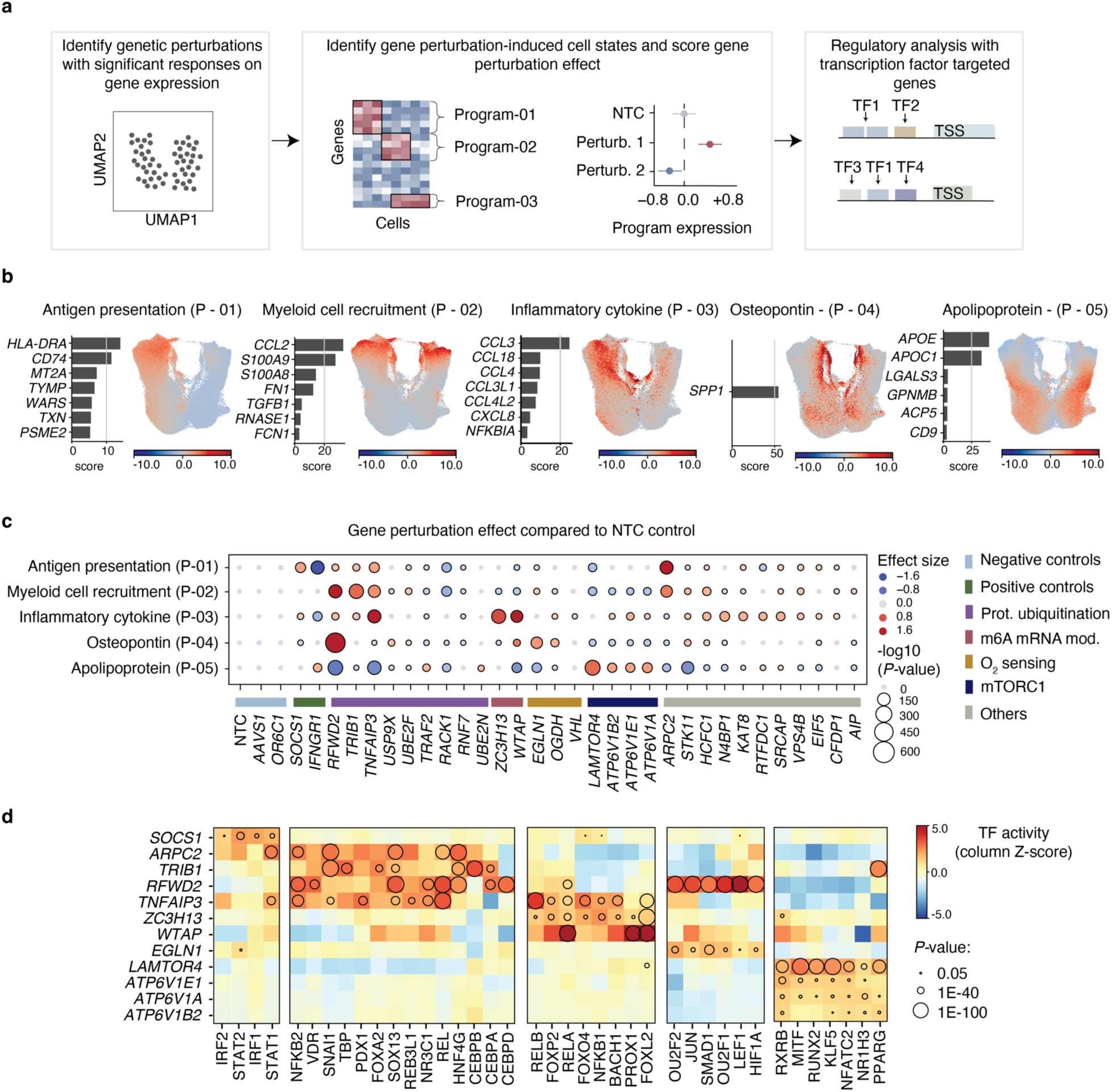
Target genes modulate a spectrum of regulatory circuits controlling macrophage states. **a,** Workflow of key analysis features. **b**, Programs altered by the introduced perturbations. Program expression scores in UMAP projection and the top correlated genes within each program. **c**, Target gene perturbation effect, indicated as effect size across each program compared to NTC controls in IFN-γ stimulated cells. Dot color represents effect size, and dot size corresponds to negative base 10 log (*P* value). **d**, Z-score normalized activity scores (color bar) of selected TFs (columns) significantly (*P* value < 0.05) induced upon perturbation of indicated target genes (rows) (full matrix in Supplementary Figure 5a).

Briefly, to identify perturbations with no significant impact on gene expression, we measured global transcriptional changes using a permuted energy distance test and compared all perturbations against the NTC cells at the level of the PCA latent space to test which cells were more likely to be drawn from the control population. Using this metric, six perturbations (*BIRC3*, *SUPT16H*, *IRF2*, *VIRMA*, *METTL14*, and *METTL3*) did not cause significant transcriptional changes after 1000 permutations (**Supplementary Fig. 3d)**. Indeed, these six perturbations were most similar to the NTC control according to cosine similarity in the same latent space (**Supplementary Fig. 3e)** and showed the least differentially expressed genes when compared to the NTC, representing potentially incomplete gene perturbation (**Supplementary Fig. 3d).** These findings precluded further analysis of the six genes. Since gene perturbations spanned a continuum of states rather than well delineated clusters, Latent Dirichlet Allocation (LDA), a probabilistic grade-of-membership model, was used to identify transcriptional programs (set of genes with co-varying expression), across the perturbation dataset (**Fig. 2b, Supplementary Fig. 4a**). To assess IFN-γ induced effects, each LDA-derived program was scored across treatment conditions **(Supplementary Fig. 4b).** Next, the effect size of each target gene perturbation for each LDA-derived program was determined by estimating how transcriptional program expression in cells from each individual perturbation deviated from the NTC control cells (**Fig. 2c, Supplementary Fig. 4c**). Finally, regulatory circuits associated with each perturbation were identified by estimating transcription factor activities using the collection of transcription factors (TF) and their transcriptional targets provided by DoRothEA. 127 TFs with significantly enriched activities across 32 perturbations were recovered (**Fig. 2d, Supplementary Fig. 5a, Methods**).

### Target genes modulate a spectrum of regulatory circuits controlling macrophage states

This model captured five programs, characterized by genes involved in different immune and cellular processes, which were annotated as: antigen presentation (*HLA-DRA* in P01), myeloid cell recruitment (*CCL2* in P02), inflammatory cytokine (*CCL3* in P03), osteopontin (*SPP1* in P04), and apolipoproteins (*APOE* and *APOC1* in P05) (**Fig. 2b and Supplementary Fig 4a**).

Perturbation of 29 target genes, previously identified in the pooled genome-wide screen, had significant effects on each of the 5 programs revealed (**Fig. 2c**). Of note, deletion of one gene can introduce transcriptional effects on other genes through various pathways with direct regulatory or indirect effects. As such, most target genes do not strictly regulate one transcriptional program, but rather affect multiple programs simultaneously. For example, *RFWD2* deletion induced a significant effect on each of the 5 programs revealed through positive or negative effects. RFWD2 (also known as COP1) is an E3 ubiquitin ligase which targets multiple proteins implicated in cell cycle regulation and differentiation but also inflammatory signaling for degradation^30^. Thus, RFWD2 has annotated roles in multiple regulatory networks; this was also reflected in our screen. To provide clarity in our model, we hereafter discuss gene perturbations with large effect sizes for each of the five programs revealed.

The antigen presentation program (P01) is characterized by canonical IFN-γ-response genes including several MHC class II components and interferon-stimulated genes (ISGs) and strongly enriches in IFN-γ primed cells (**Fig. 2b, Supplementary Fig 4b**). As expected for known regulators of JAK/STAT signaling, deletion of the IFN-γ receptor, *IFNGR1,* led to downregulation of P01, while *SOCS1* deletion, a major negative regulator of STAT/JAK signaling downstream of IFN-γ, upregulated P01 (**Fig. 2c**). Our TF analysis confirmed increased activity of STAT1/IRF1 upon deletion of *SOCS1* (**Fig. 2d**). Thus, the detection of established links between key regulators of IFN-γ response pathway and regulated TFs confirms the fidelity of our perturbation and analysis approach.

Besides the established role of *SOCS1,* the actin filament nucleation protein *ARPC2*, was identified as a novel negative regulator of P01 (**Fig. 2c**). Interestingly, deletion of *ARPC2* increased TF activity of STAT1 to a similar degree as *SOCS1* (**Fig. 2d**). Polymorphisms in *ARPC2* along with MHC class II and components of the STAT1 signaling pathway have been described as susceptibility genes for ulcerative colitis^31^, thus highlighting a potential clinical relevance of this regulatory circuit.

The myeloid cell recruitment program (P02) is characterized by genes impacting cell migration including *CCL2* and calprotectin (*S100A8/9)* (**Fig. 2b**). Macrophage states associated with P02 genes have been identified in disease indications associated with high mononuclear cell infiltration, such as rheumatoid arthritis (RA) and cancer^32,33^. Multiple perturbations led to significant upregulation of P02, among them components of E3 complexes, RFWD2 and TRIB1, which control the turnover of AP-1 and CEBP family members through ubiquitination^34–36^ (**Fig. 2c**). *RFWD2* deletion enhanced TF activity of CEBPA, CEBPD and JUN while *TRIB1* deletion enhanced CEBPA and CEBPB among others in hMDMs (**Fig. 2d**). In line with these findings, upregulation of AP-1 TFs were found in synovial tissues along with an increase of CCL2 serum levels in RA patients^37^. These results are consistent with previous findings suggesting that *RFWD2* and *TRIB1* control a macrophage state which promotes myeloid cell recruitment by regulating the turnover of AP-1 and CEBP family members.

The inflammatory cytokine program (P03) is characterized by CC-family mediators including *CCL3*, *CCL18* and *CCL4* (**Fig. 2b**). Deletion of the central negative regulator of NFκB signaling *TNFAIP3* led to significant upregulation of P03 and elevated activity of NFκB1 and RELA, as expected (**Figs. 2c, d**). In addition to *TNFAIP3*, deletion of *ZC3H13* and *WTAP*, two components of the m6A writer complex, led to upregulation of P03 as well as elevated activity of RELA, thus suggesting a regulatory role of m6A on NFκB signaling in human macrophage biology (**Figs. 2c, d**).

The osteopontin program (P04) is defined by *SPP1* expression (**Fig. 2b)**. Osteopontin (encoded by SPP1) is highly upregulated in macrophages associated with fibrosis and cancer^4,38–40^ and associated with poor clinical outcome in colorectal cancer^4^. Deletion of the well-studied negative regulator of hypoxia *EGLN1* (encoding PHD2) led to upregulation of P04. Indeed, SPP1 is highly induced in hMDMs cultured in hypoxic conditions, suggesting that P04 represents a hypoxic macrophage state (**Supplementary Fig. 5b**). In addition to *EGLN1,* our screen highlights a regulatory role of RFWD2 on P04 (**Fig. 2c**). The activity of multiple TFs including the master transcriptional regulator of hypoxia HIF1A were elevated in *EGLN1* and *RFWD2* deleted cells (**Fig. 2d**). A functional role of RFWD2 on hypoxia has no prior literature evidence, warranting focused investigation in future studies.

The apolipoprotein program (P05) is characterized by apolipoprotein coding genes *APOE* and *APOC1,* among others (**Fig. 2b**). Deleting upstream regulators of the mTORC1 signaling pathway including components of the v-ATPase (e.g., *ATP6V1B1*) and *LAMTOR4,* a component of the Ragulator complex, led to upregulation of P05 (**Fig. 2c**). Concomitantly, a larger set of TFs involved in the regulation of nutrient metabolism including PPARG and NR1H3 showed enhanced activity in this group of perturbations (**Fig. 2d**). In clinical settings, apolipoprotein expressing macrophages were identified in the tumor environment in breast and colorectal cancer patients^7,41^. These findings suggest a role of the apolipoprotein-associated macrophage state in dysregulated nutrient or growth factor sensing through mTORC1 inhibition.

Additional programs characterized by macrophage-centered genes including *MMP9* (P07) or *LYZ* (P10) were revealed in this study (**Supplementary Fig. 4a**) further adding to the spectrum of cell states and their regulatory circuits. Together, transcriptional analysis of target gene perturbations in hMDMs revealed a spectrum of cell states relevant to human disease, and discovered intrinsic regulators of these states.

### Target genes promote distinct cytokine secretion profiles

In addition to their impact on macrophage transcriptional states, we hypothesized that the identified genes would also affect macrophage cytokine secretion profiles. To test this directly, the top 7 target genes with the highest effect size (size > 0.7) on the perturbation-induced programs P01-05 (**Fig. 2c**) were deleted, followed by quantification of secreted inflammatory chemokines and cytokines (**Figs. 3a, b** and **Supplementary Fig. 6a**). For reference, positive controls (*SOCS1* and *IFNGR1*) and a negative control (NTC) were also included.

**Fig. 3:**
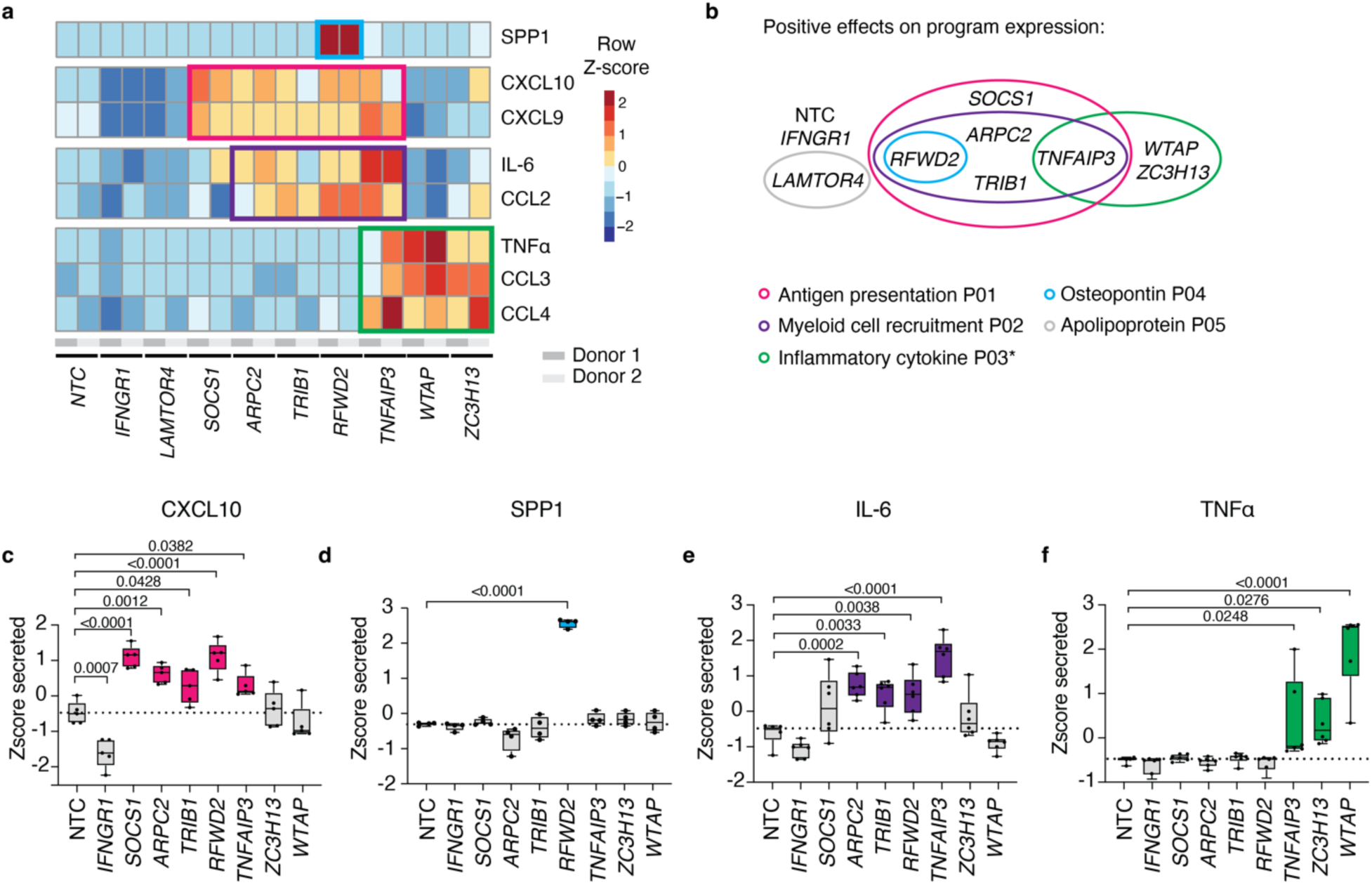
Selected target genes promote distinct cytokine secretion profiles. **a,** Heatmap of secreted cytokine and chemokine measurements upon IFN-γ priming. Z-score scaling shown for two individual donors (full matrix in Supplementary Figure 6a) **b**, Positive effect sizes on each of the 5 programs by selected perturbations. For all programs, grouping of perturbations is based on significance, but for P03 (*), only perturbations with effect sizes > 0.8 are included. **c-f**, Secreted levels of CXCL10, SPP1, IL-6 and TNFα indicated as Z-score for each cytokine. Points represent a single gene perturbation and donor measurement (CXCL10, IL-6, TNFα n=5; SPP1 n=4). Summary data are shown as mean, with *P* values determined by ANOVA with Dunnett’s multiple comparisons test.

Deletion of selected target genes caused measurable changes in chemokine and cytokine secretion levels in two independent donors, with the exception of *LAMTOR4* (**Fig. 3a**). Interestingly, target gene perturbations clustered according to the macrophage transcriptional programs they impacted. To further validate this, additional perturbation experiments were performed in 5 independent donors, examining one representative cytokine for each group (**Figs. 3c-f**).

Perturbations with significant positive effect sizes on the IFN-γ-driven antigen presentation program P01, denoted as P01 regulators, enhanced the canonical IFN-γ-induced cytokines CXCL9 and CXCL10 (**Figs. 3a, c**). Similarly, deletion of *RFWD2,* a regulator of the Osteopontin program P04, induced elevated secretion of SPP1 itself, in line with *SPP1* being the top scoring gene for P04 (**Figs. 3a, d**). Interestingly, deleting regulators of the myeloid cell migration program P02 enhanced both CCL2 and IL-6 secretion (**Figs. 3b, e**). While *CCL2* was the top scoring gene for this program, IL-6 transcript was not associated with any of the identified programs, despite multiple IL-6 promoting cis-regulatory elements such as CEBPB, AP1 family members and NFκB^42^ being upregulated in P02 regulator perturbed cells (**Fig. 2d**). Cytokine secretion profiling thus provided additional insights into macrophage effector functions such as IL-6 secretion that align with the regulatory circuitry revealed in our scCRISPR screen but were not captured by transcriptional profiling.

Deleting regulators of the inflammatory cytokine program P03 including *TNFAIP3*, *ZC3H13* and *WTAP* induced a distinct cytokine secretion profile characterized by elevated levels of CCL3, CCL4 and TNFα (**Figs. 3a, f**). Strikingly, TNFα transcript levels were barely detectable in our perturbation dataset despite the ∼ 30-fold elevated TNFα protein levels detected in *WTAP* deleted cells, highlighting again the utility of combining transcriptional profiling with cytokine secretion profiling (**Supplementary Fig. 6b**).

Given that the scCRISPR screen indicated a regulatory role for NFκB signaling underlying P03, we hypothesized TNFα to be the causal driver of this program. We tested this hypothesis focusing on the understudied regulators *ZC3H13* and *WTAP*. Pharmacological inhibition of TNFα protein by TNFRII-Fc in *WTAP-* and *ZC3H13*-perturbed cells significantly reduced P03’s top scoring genes *CCL3*, *CCL4* and *CXCL8* (**Supplementary Fig. 6c**). Additionally, hMDMs co-stimulated with both IFN-γ and TNFα displayed elevated P03 expression compared to hMDMs stimulated with IFN-γ alone (**Supplementary Fig. 6d**). Thus, loss of WTAP and ZC3H13 promoted expression of program P03 via overproduction of TNFα.

### WTAP associates with the TNFα-driven inflammatory macrophage state P03 across disease conditions

Chronically activated immune cells commonly upregulate checkpoint proteins as negative feedback loops to prevent aberrant pro-inflammatory signaling that exacerbates inflammation. Given the above demonstration of a regulatory role of *ZC3H13* and *WTAP* in TNF production, we hypothesized that these may also act as checkpoints in TNFα-induced inflammation and be upregulated under these conditions.

We selected 5 previously published scRNA-seq datasets of myeloid-rich inflammatory disease-affected tissues, and observed an enrichment of P03 in the macrophage compartment when compared to the respective healthy or uninflamed control tissues (**Supplementary Fig. 7a**). Pathway analysis demonstrated significant enrichment of TNFα and NFκB signaling in P03-high macrophages (**Supplementary Fig. 7b**). Thus, P03 represents an established TNFα-driven pathological macrophage state across disease conditions.

Next, we assessed expression levels of *WTAP* and *ZC3H13* within the selected datasets. While *ZC3H13* transcripts were inconsistent or barely detected, elevated levels of *WTAP* were consistently observed in the macrophage compartment within inflammatory disease when compared to the respective healthy or uninflamed control tissues (**Supplementary Fig. 7a**). This association was further confirmed by *in situ* hybridization (ISH) in ulcerative colitis compared with healthy colon samples. Specifically, elevated *WTAP* transcript co-localized with the macrophage (*CD68*+) compartment of inflamed tissues as compared to healthy controls. Furthermore, *WTAP* was also enriched in *CCL3*+ cells, a top marker gene of P03, indicating its association with an ongoing TNFα-induced inflammatory response (**Supplementary Figs. 7c, d**). Thus, *WTAP* is elevated in inflammatory myeloid cells across disease conditions. Together with the previous genetic perturbation data, these findings suggest a regulatory mechanism to counteract TNFα-driven inflammation.

### *ZC3H13-* or *WTAP*-perturbed hMDMs promote TNFα-dependent paracrine effects

To assess possible paracrine effects of *ZC3H13*- or *WTAP*-perturbed macrophage states, we set out to evaluate the impact of macrophage-derived cytokines on core components of the macrophage’s natural milieu in the intestine. Here, endothelial and intestinal epithelial cells were chosen to represent this milieu, since macrophages closely interact with these cell types under homeostatic conditions in the gut^43^. Exposing human endothelial cells derived from normal vasculature to conditioned media from *ZC3H13*- or *WTAP*-perturbed hMDMs upregulated the inflammation-mediated activation markers ICAM-1 and VCAM-1 in a TNFα-dependent manner (**Supplementary Figs. 8a, b**).

To assess if *ZC3H13*- or *WTAP*-perturbed hMDMs affected intestinal epithelial barrier integrity, we established human intestinal organoid monolayers derived from intestinal crypts isolated from healthy donors and exposed these to conditioned media from IFN-γ-stimulated *ZC3H13*- or *WTAP*-perturbed hMDMs (**Supplementary Fig. 8c**). Such treatment damaged the epithelial monolayer, as evidenced by transfer of cell-impermeable dextran across the monolayer (**Supplementary Fig. 8d**). Pharmacological inhibition of TNFα by TNFRII-Fc rescued barrier leakage (**Supplementary Fig. 8d**). Consistent with these findings, epithelial monolayer viability was decreased upon exposure to conditioned media from stimulated *ZC3H13*- or *WTAP*-perturbed hMDMs and rescued by TNFRII-Fc treatment (**Supplementary Fig. 8e**). Thus, our data thus indicates that *ZC3H13*- or *WTAP*-perturbed macrophages promote paracrine effects in *in vitro* environments, in addition to the autocrine effects observed in hMDMs.

### Perturbation of the m6A writer complex elevates TNF**α**

Together with other protein components such as METTL3 and METTL14, WTAP and ZC3H13 form the “m6A writer complex”, which co-transcriptionally mediates mRNA methylation^27,28^. The main function of m6A, the most abundant internal mRNA modification, is the regulation of mRNA fate, in particular by promoting degradation of modified molecules, thereby regulating cell development, differentiation, and other processes^27,28^.

Intriguingly, despite being revealed as target genes in our genome-wide screen, *METTL3-* and *METTL14*-perturbations in hMDMs did not significantly change gene expression (**Supplementary Figs. 3d, e**). Previous studies have reported the expression of alternatively spliced *METTL3* transcript isoforms which can bypass CRISPR/Cas9 mutations to express functionally active methyltransferase^44^. We verified that a short *METTL3* isoform was indeed expressed in *METTL3-*perturbed hMDMs but not NTC control cells (**Supplementary Fig. 9a)**. Using an improved sgRNA guide design targeting either exon 5 or 6, which are part of the coding sequence for the methyltransferase domain, ablated the full length *METTL3* isoform as well as the appearance of the short *METTL3* isoform in hMDMs and confirmed elevated levels of TNFα secretion as well as elevated P03 expression (**Supplementary Figs. 9b, c, d)**. Additionally, we designed an improved sgRNA guide for METTL14 to achieve highly efficient protein depletion, and in doing so, also confirmed elevated levels of TNFα (**Supplementary Figs. 9e, f)**. Thus, efficient perturbation of WTAP, ZC3H13, METTL3 or METTL14 confirmed that loss of the essential components of the m6A writer complex elevates TNFα secretion in IFN-γ primed hMDMs. While dysregulation of m6A has been implicated in various inflammatory diseases and cancer^45^, how m6A regulates macrophage function and cytokine response is poorly understood, with inconclusive or contradictory findings^46–52^, prompting us to further investigate the role of m6A on TNFα in hMDMs.

### WTAP and ZC3H13 control m6A modification of human TNF mRNA

Multiple other stimuli beyond IFN-γ, such as microbial ligands, can elicit TNFα. To understand if m6A broadly regulates TNFα production, TNFα cytokine release in *WTAP*- or *ZC3H13*-perturbed hMDMs was analyzed upon TLR3 activation with poly I:C (a substitute for viral dsRNA), TLR7/8 activation with R848 (Resiquimod), or TLR4 activation with LPS. Perturbation of *WTAP* or *ZC3H13* substantially elevated P03 expression and TNFα cytokine release following stimulation with all three TLR ligands (**Fig. 4a, Supplementary Fig. 10a**), demonstrating that m6A is a fundamental, context-independent regulator of TNFα.

**Fig. 4:**
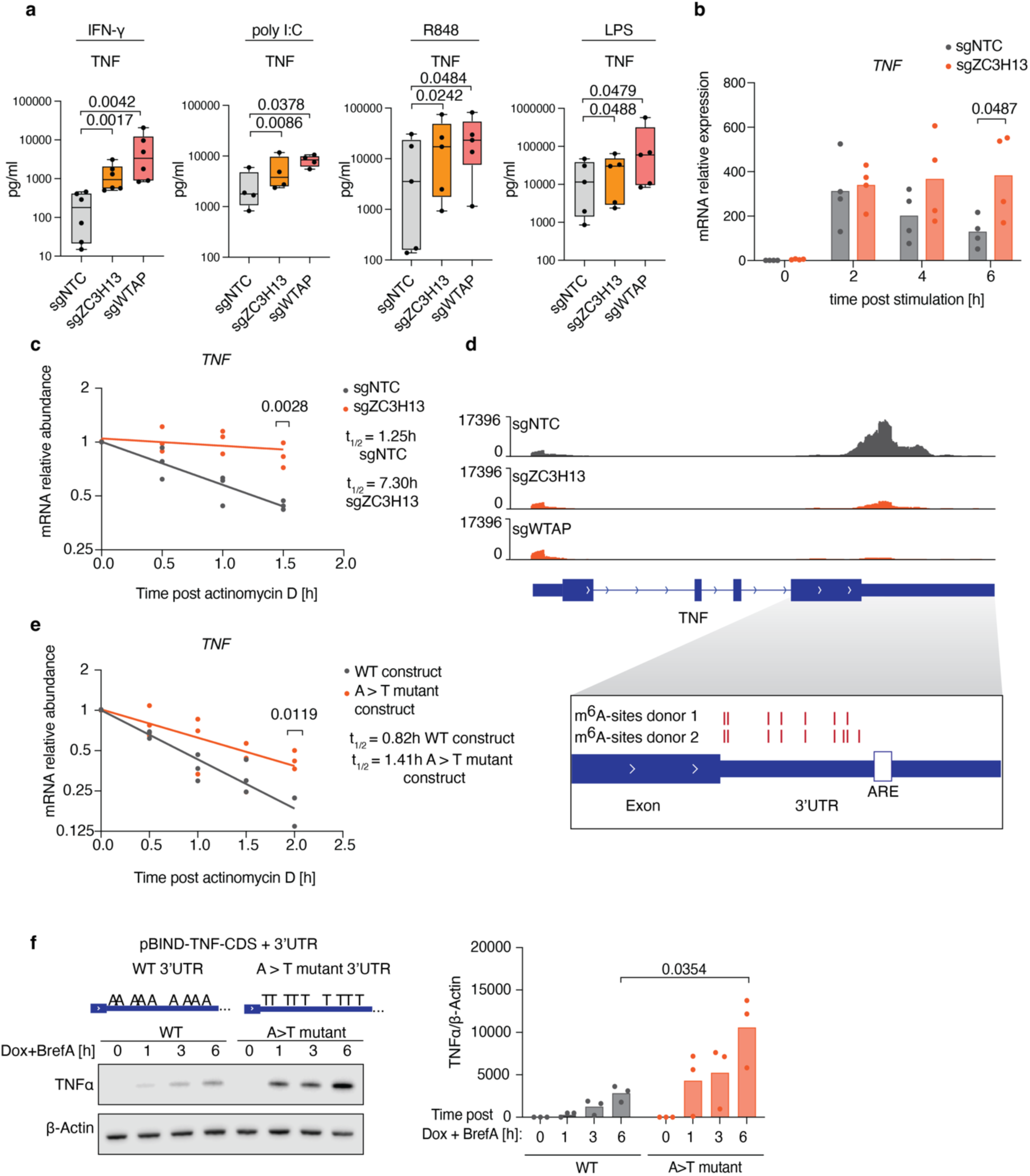
*TNF* mRNA is m6A modified. **a**, TNFα in the culture medium of genetically perturbed hMDMs 18 h after indicated stimulations. **b**, Relative *TNF* mRNA levels and **c**, half-life measurements upon Actinomycin D treatment in *ZC3H13*-perturbed and NTC control hMDMs. **d**, Genomic visualization of normalized m6A-eCLIP signal along the human *TNF* transcript in NTC control, *WTAP-* and *ZC3H13*-perturbed hMDMs. m6A residues indicated as red lines as identified in two individual donors, ARE = AU-rich element. **e**, *TNF* mRNA half-life measurements using a reporter construct (coding region of *TNF* fused to either a wildtype 3’UTR (WT) or a 3’UTR in which the 9 m6A-modified adenosines were mutated to thymidine (A>T mutant)) upon Actinomycin D treatment. **f**, TNFα expression levels induced by A>T mutant construct described in (e). (a, b, f) Data represents mean values. a, b, c, each symbol represents an individual donor (a, n= 4 - 6), (b, n=4), (c, n=3). Data are derived from 2 (b) and 3 (a, c, e, f) independent experiments. *P* values determined by Students two-sided t-test.

Given that loss of WTAP or ZC3H13 increased TNFα levels irrespective of the nature of the pro-inflammatory stimulus, we wondered whether m6A acted directly on the *TNF* transcript. Since *WTAP*-depletion compromised the basal fitness of cells, we focused on *ZC3H13*-depletion for these mechanistic studies. Monitoring *TNF* transcript levels over time revealed sustained levels in *ZC3H13*-depleted but not in control hMDMs upon stimulation (**Fig. 4b**). Similarly, we observed enhanced *TNF* transcript stability following transcriptional inhibition by Actinomycin D in *ZC3H13*-depleted, but not in control hMDMs. In contrast, *IFNB1* and *IL6* transcript levels were not significantly affected (**Fig. 4c, Supplementary Fig. 10b, c**). Thus, m6A likely affects the stability of a subset of inflammatory mediators such as *TNF* instead of acting globally on cytokine transcripts.

Seeking a mechanistic understanding of *TNF* transcript regulation by m6A, m6A-enhanced crosslinking and immunoprecipitation (m6A-eCLIP) was performed to profile site-specific m6A modifications at the transcriptome level. The majority of high-confidence m6A peaks mapped to the coding regions and three prime untranslated regions (3’UTRs) containing the DRACH (D=A, G or U; R=A or G; H = A, C or U) sequencing motif^53,54^ (**Supplementary Fig. 10d**). Deletion of *WTAP* or *ZC3H13* induced strong reductions in global m6A levels (**Supplementary Fig. 10d**). Importantly, the m6A-eCLIP analysis revealed multiple m6A peaks mapping to *TNF* mRNA (**Fig. 4d, Supplementary Fig. 10e**). Further analysis identified a total of nine methylated adenosines specifically located at the 3’UTR of *TNF*, of which seven were conserved across both donors (**Fig. 4d**). Deletion of *WTAP* or *ZC3H13* drastically reduced m6A peaks on *TNF* transcript in both donors, demonstrating that these m6A writer components are required for *TNF* m6A modification in hMDMs (**Fig. 4d, Supplementary Fig. 10e**). Further analysis of *IFNB1* and *IL6* transcripts demonstrated remarkably reduced peak abundance and sites as compared to *TNF* (**Supplementary Figs. 10f, g)**, consistent with the lack of impact on *IFNB1* and *IL6* mRNA stability in *ZC3H13*-perturbed hMDMs (**Supplementary Figs 10h**).

To functionally demonstrate that impaired m6A of *TNF* affects transcript stability, a reporter system was designed in which all m6A modification sites on *TNF* were mutated (A>T mutant). Expression of the A>T mutant construct yielded increased *TNF* transcript stability and markedly elevated TNFα protein levels (**Figs. 4e, f**). These results confirm the role of m6A methylation sites in regulating *TNF* transcript stability and in turn TNFα protein levels. Additionally, deletion of *YTHDF2*, an m6A reader protein which selectively targets m6A-modified mRNAs for degradation^55^, yielded a significant increase in TNFα levels, suggesting that m6A of *TNF* transcript acts as an mRNA degradation signal (**Supplementary Fig. 6d**).

### Two SNPs predicted to impact m6A installation at the 3’UTR of *TNF* associate with cystic kidney disease

Phenome-wide association studies (PheWAS) provide an opportunity to identify genetic variants or a collection thereof linked to human disease by scanning thousands of human phenotypes. We performed a PheWAS on SNPs localized within the DRACH sequence motif of identified m6A sites on *TNF* 3’UTR, thus predicted to impact m6A installation. The PheWAS was performed on individuals using the UK Biobank whole-genome sequencing (WGS) data, as the majority of these SNPs were not detected in the UK Biobank imputed array or whole-exome sequencing (WES) data, suggesting that these non-coding variants are not well covered. Thus, a collapsing analysis using WGS data provided a better opportunity to identify rare non-coding variants linked to human disease. While the necessity for WGS data restricted our PheWAS to the UK Biobank, currently the only resource which couples WGS with phenotypic data, it still allowed for a comprehensive analysis as it represents the largest collection of WGS on nearly 500,000 individuals^56^.

Our analysis revealed an association with cystic kidney disease, an inflammatory pathology known to be driven by TNFα^20–22^ (**Figure 5a)**. Specifically, identified carriers harbored one of the two SNPs associated with disease (rs190947828 or rs777492465) (**Figure 5b)**. Of note, P03 was highly elevated in a previously published dataset of polycystic kidney disease and highly enriched in the macrophage compartment^57^ (**Figure 5c-e)**. These novel associations implicate non-coding germline variation in *TNF* as relevant contributors to inflammatory pathology and prompt further investigation as to why these observed associations emerge in a disease-specific manner.

**Figure 5:**
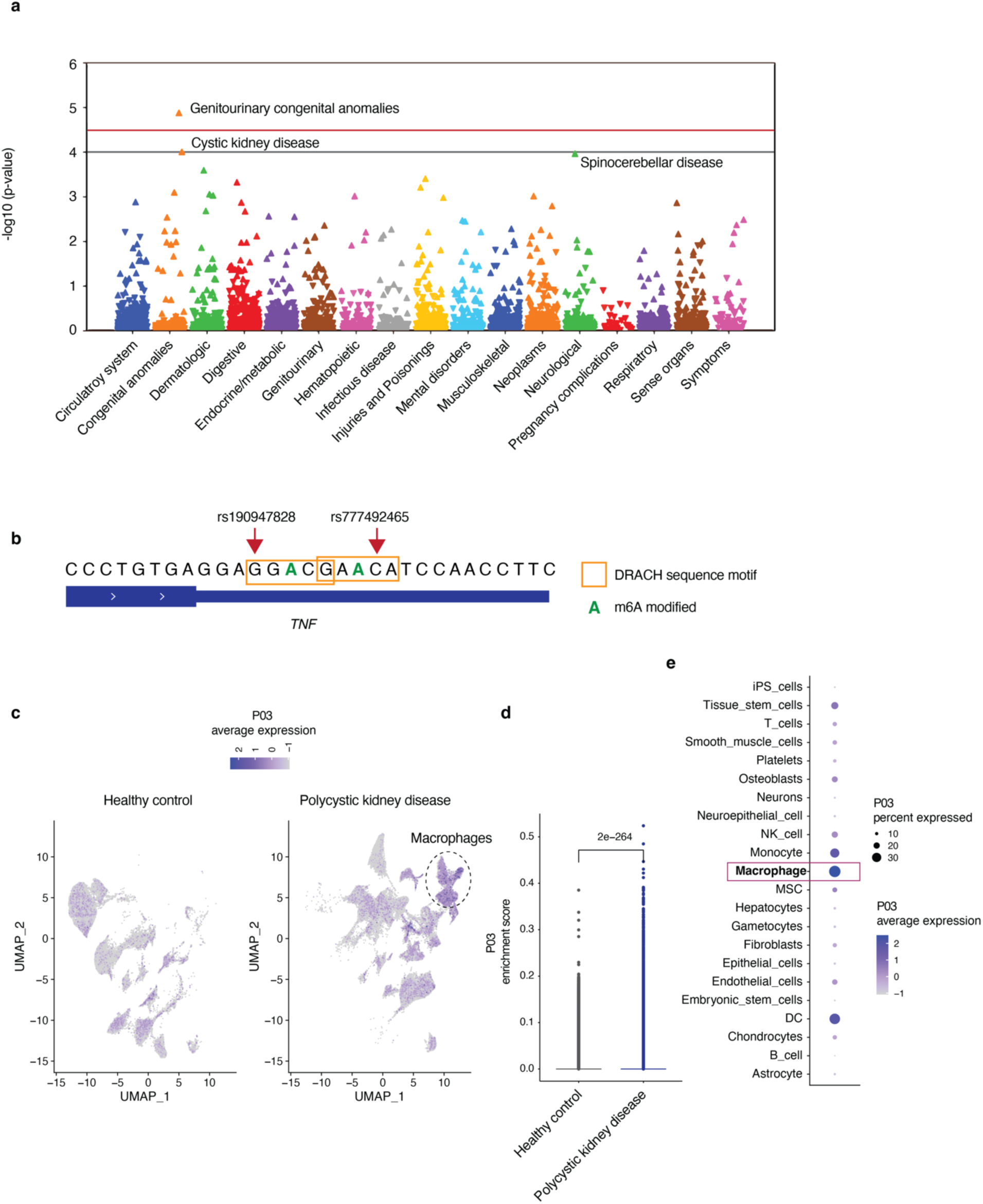
PheWAS analysis of genomic locations predicted to affect m6A installation on *TNF* mRNA. **a,** Manhattan plot indicating PheWAS associations between the aggregated set of SNPs (predicted to impact m6A installation on *TNF* mRNA) and 1,695 phecodes (binary traits) using 424,136 UKB individuals. The red line indicates Bonferroni correction (-log10(<0.05/# of phecodes>)) while the gray line indicates a suggestive threshold based on previous work^60^. Associations [-log10(*P* value)] are represented as triangles with the direction facing up/down indicating increased/decreased risk respectively in carriers relative to non-carriers. **b**, Genomic visualization of the two SNPs associated with cystic kidney disease along the *TNF* transcript. **c**, UMAP showing average expression of P03 and **d**, P03 enrichment scores in healthy kidney and polycystic kidney disease^57^. Box plots summarize the median, interquartile and 75% quartile range for each tissue type. *P* values determined by two-sided t-test **e**, P03 expression across each cell type in polycystic kidney disease.

## DISCUSSION

High resolution single-cell gene expression profiling has revealed pro-inflammatory macrophage states across numerous human pathologies^5^, but our understanding of the underlying regulatory circuits governing these states is limited. This is in part due to technical challenges hampering the use of large-scale CRISPR-Cas9 mediated screening approaches on primary human macrophages, as well as potentially divergent biology of myeloid cells in model organisms. The resulting knowledge gap limits our ability to identify regulatory mechanisms underlying inflammatory macrophage states that are associated with disease and to reveal novel, actionable targets for therapeutic intervention.

To overcome these limitations and enable the systematic discovery of intrinsic regulators of inflammatory macrophage function, we developed a funneled CRISPR screening approach across species. This allowed for unbiased, genome-wide screening in primary murine macrophages followed by targeted interrogation of the most potent perturbations in human primary cells. Using transcriptional profiling in hMDMs, we resolved five distinct macrophage cell states relevant to human disease and discovered intrinsic regulators of these states *de novo*. Multiplexed secretome analysis revealed that these states are associated with distinct cytokine secretion profiles. Of note, key inflammatory cytokines such as IL-6 and TNFα were not captured at the transcript level in any of the identified programs, possibly due to transient or low expression levels, but were revealed at the protein level, highlighting the power of a multimodal approach.

The screen captured distinct programs associated with key features of macrophage function such as antigen presentation, myeloid cell recruitment, inflammatory cytokine expression, osteopontin secretion and apolipoprotein release, revealing a breadth of roles for the identified target genes, several of which are novel. These include ARPC2 and its role in restricting a STAT1/IRF1 mediated antigen presentation program, RFWD2 and regulation of a hypoxic state characterized by SPP1 expression, mTORC1 signaling components and the apolipoprotein state, and m6A writer components WTAP, ZC3H13 and METTL3 in suppressing a TNFα-driven inflammatory state.

Well-known negative regulators of TNFα-mediated inflammation, such as A20 encoded by *TNFAIP3*, fine-tune the magnitude and length of a TNFα-driven cell state. Here, we describe m6A modification as a novel mechanism for restricting a TNFα-driven state in human macrophages, where m6A modification of specific adenosines in the 3’UTR of *TNF* mRNA by WTAP and ZC3H13 promotes destabilization of *TNF* and thereby restricts TNFα cytokine levels. While m6A likely regulates additional mechanisms beyond *TNF* transcript stability that may also participate in tuning TNFα, our findings show that *TNF* m6A modification controls *TNF* transcript decay rate and thereby affects TNFα secretion levels, which in turn control a TNFα-driven cell state in human primary macrophages (**Supplementary Fig. 11**). SNPs predicted to affect m6A installation at 3’UTR of human *TNF* revealed an association with cystic kidney disease, thus implicating an m6A-mediated regulatory mechanism in human disease. Future studies on the causal roles of the two identified SNPs are needed to resolve the molecular mechanisms underlying these variants.

Dysregulation of m6A has been implicated in various inflammatory diseases and cancer^45,58,59^, but studies of how m6A regulates macrophage function appear inconclusive or contradictory and have been limited to mouse models or cell lines^46–52^. For instance, myeloid-specific deletion of METTL3 in a murine tumor model was proposed to promote expression of both pro- and anti-inflammatory cytokines by controlling *Spred2* mRNA translation^46^. A separate study suggested that loss of METTL3 in murine macrophages suppressed expression of pro-inflammatory cytokines by regulating *Irakm* transcript stability^50^. Of note, *TNF* mRNA stability was shown to not be affected by m6A in a murine macrophage cell line^50^. In contrast, our data demonstrates a role for m6A in regulating *TNF* by direct perturbation of multiple key components of the mRNA m6A machinery in primary human macrophages. Future work will be required to delineate if m6A controls macrophage phenotypes in species- or cell lineage-specific manners.

Taken together, our work demonstrates a comprehensive platform to uncover regulatory mechanisms controlling clinically meaningful macrophage states. It also highlights the importance of focusing on primary human systems to uncover translationally relevant mechanisms of pathway modulation. The funneled approach described herein can be broadly applied to investigate other macrophage phenotypes of interest and hopefully reveal key regulators of cell states beyond the ones described in this study, thereby providing novel leads for future therapeutic intervention.

## SUPPLEMENTARY FIGURES

**Supplementary Fig. 1:**
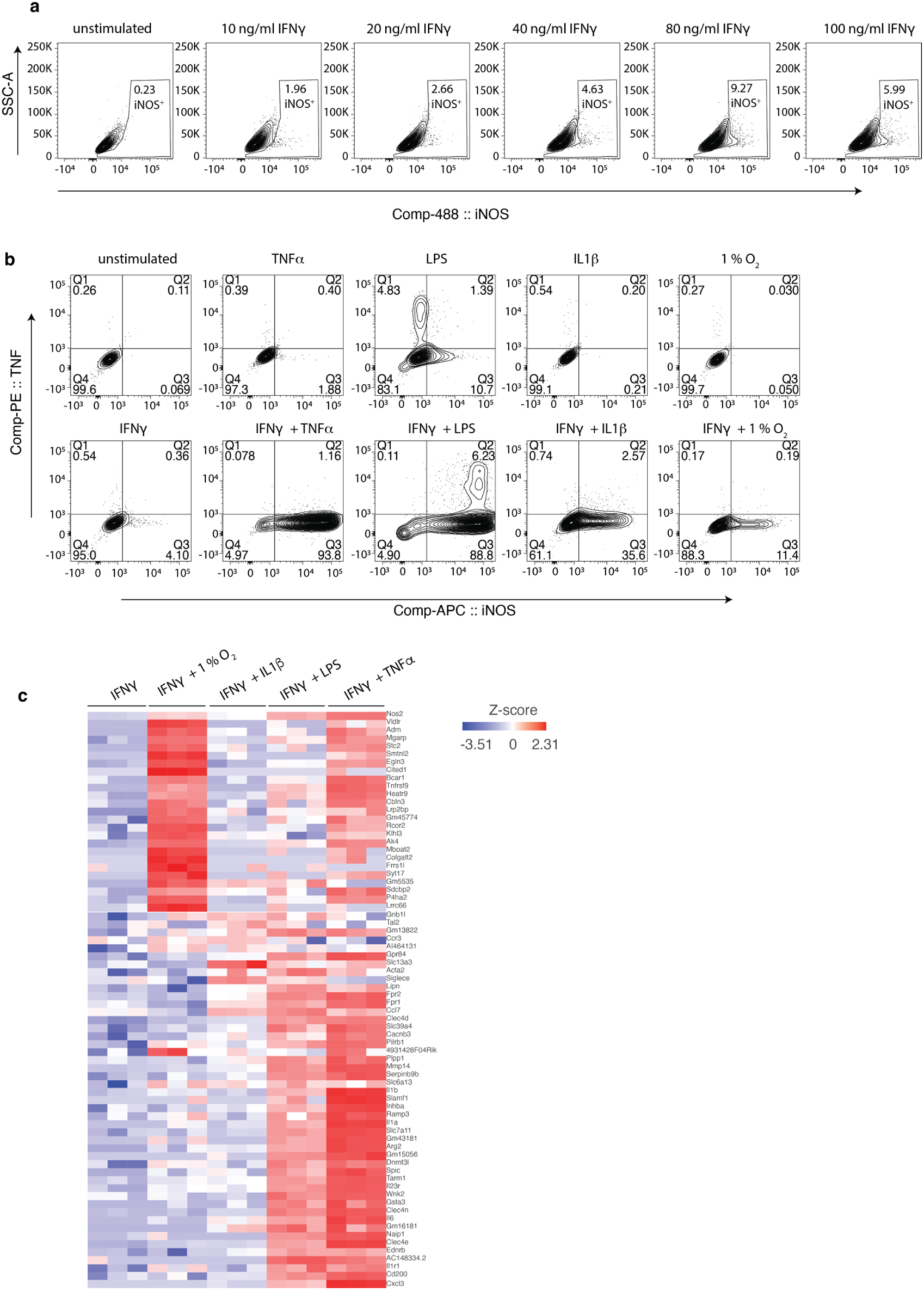
Screening design in BMDMs. **a**, Flow cytometry analysis of mBMDMs primed with IFN-γ in indicated concentrations. **b**, Flow cytometry analysis of intracellular TNFα and iNOS expression in mBMDMs stimulated as indicated. **c**, Heatmap of differentially expressed genes between each of the four comparisons in mBMDMs across 3 mice.

**Supplementary Fig. 2:**
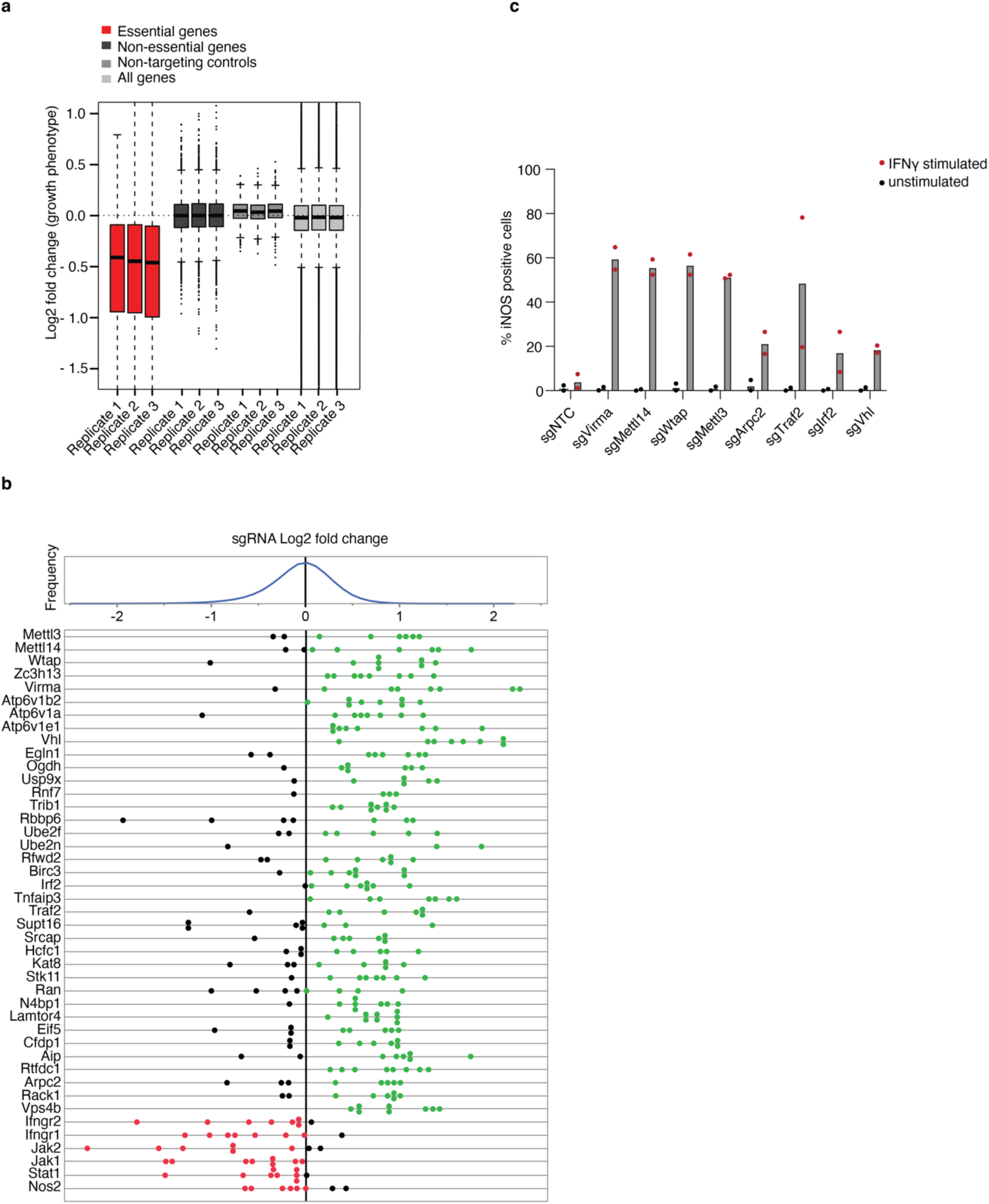
Quality controls for genome-wide CRISPR screen in mBMDMs. **a**, sgRNA drop out of essential genes across 3 individual screens (replicates 1 - 3). **b**, gRNA-level log2-fold changes of representative genes. Vertical dotted gray line represents the median of control sgRNAs. Red and blue lines indicate enriched or depleted sgRNA rank for each indicated gene respectively. **c**, Flow cytometry of intracellular iNOS expression levels in perturbed mBMDMs as indicated (n = 2).

**Supplementary Fig. 3:**
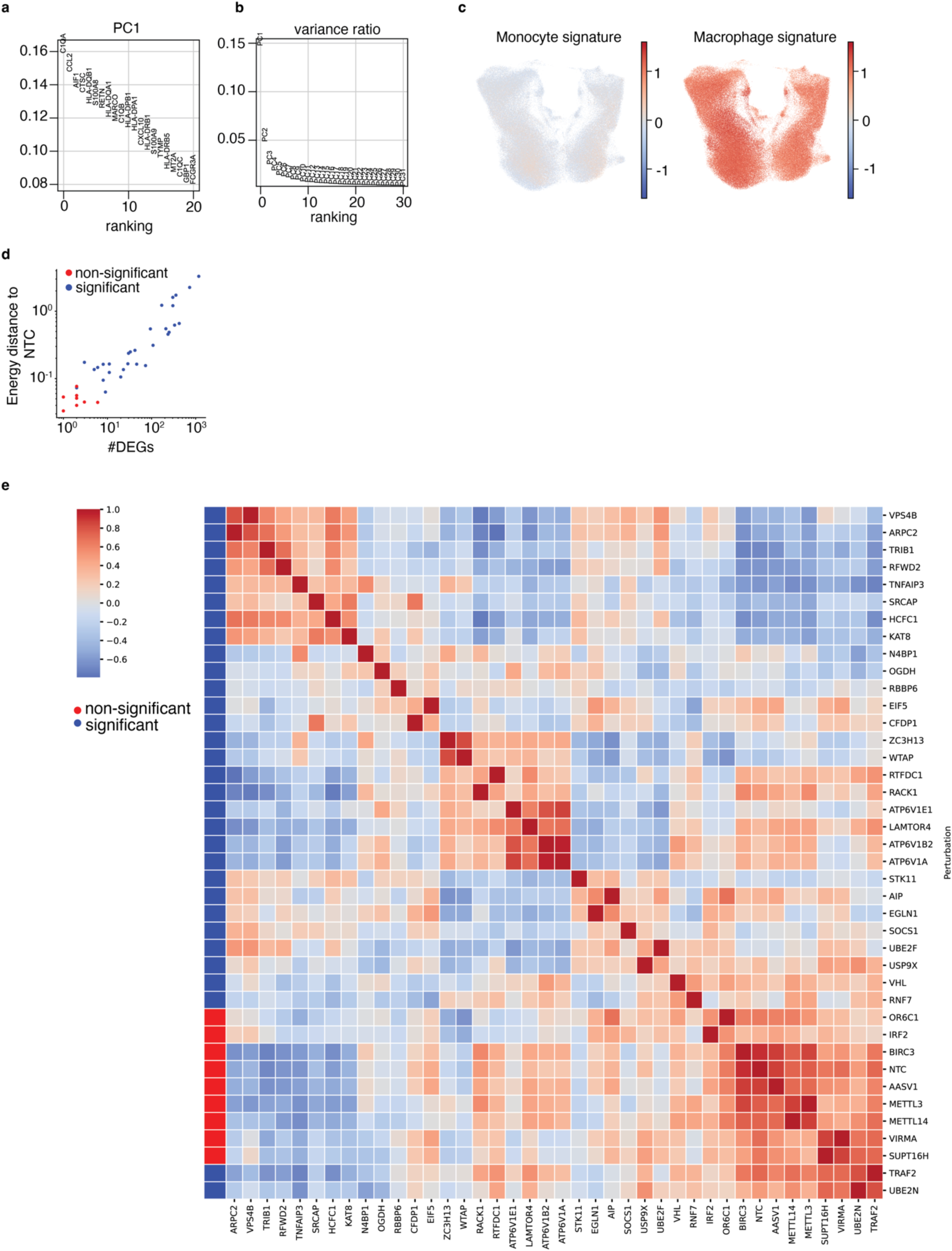
PCA and permuted energy distance test. **a,** Individual gene ranking of PC1. **b,** variance ratio of PCA. **c**, *Bona-fide* monocyte and macrophage signature score in UMAP projection. **d**, scatter plot of the energy distance against the number of differentially expressed genes (DEG) calculated as pairwise comparison to NTC. Each dot represents a perturbation. Perturbations with significant effects (energy distance; FDR < 0.01, DEGs; FDR < 0.05) indicated in blue, non-significant in red. **e**, cluster map of cosine similarity between perturbations. Rows and columns are hierarchically clustered. Perturbations with significant effects as determined in (d) indicated in blue, non-significant in red.

**Supplementary Fig. 4:**
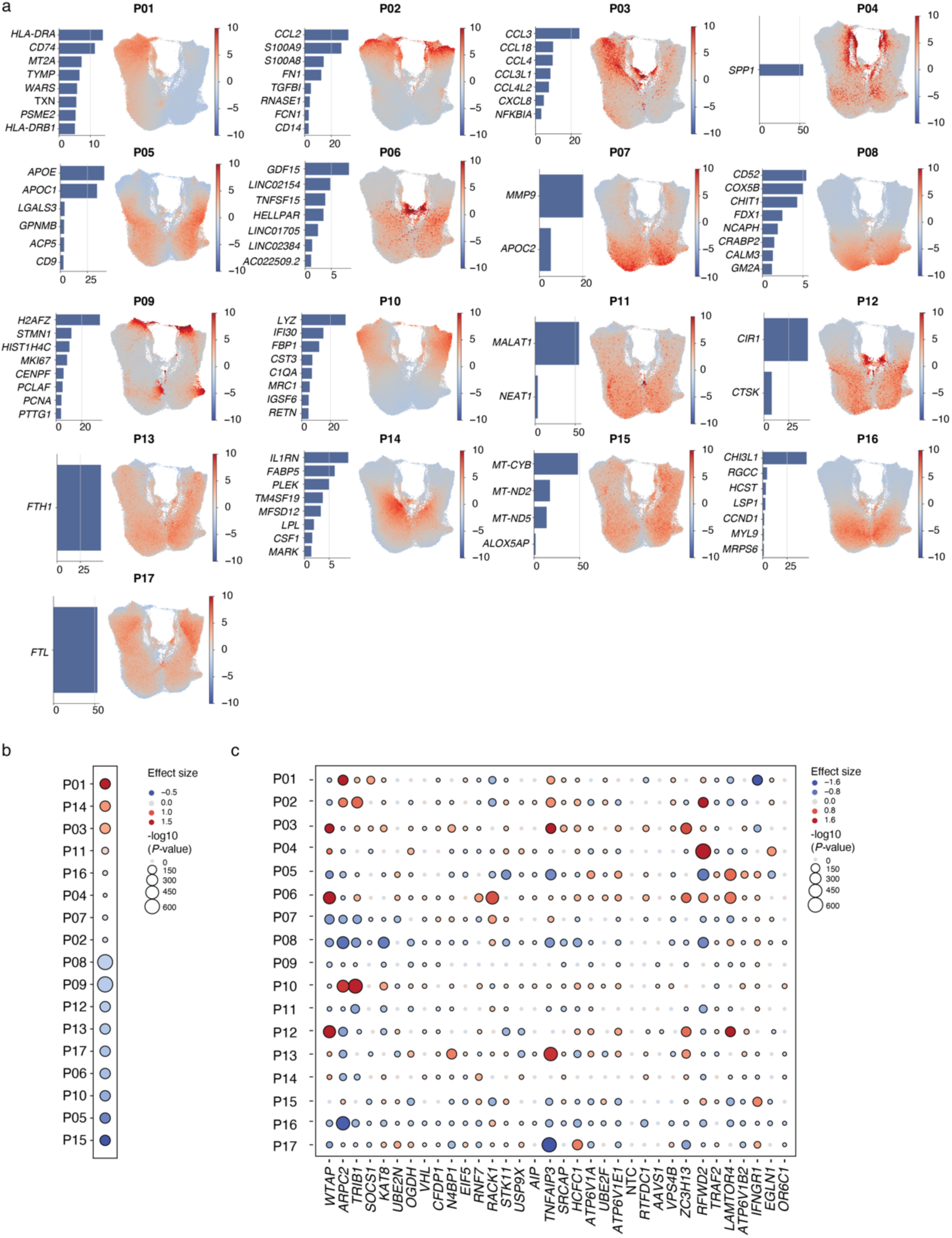
LDA-derived programs and effect sizes across all perturbations. **a**, LDA-derived programs. Feature plot of program expression scores and the top correlated genes within each program. **b,** Program enrichment upon IFN-γ priming. **c,** Genetic perturbation effects across programs compared to NTC control in IFN-γ primed conditions. b, c, Dot color represents effect size, and dot size corresponds to negative base 10 log (*P* value).

**Supplementary Fig 5:**
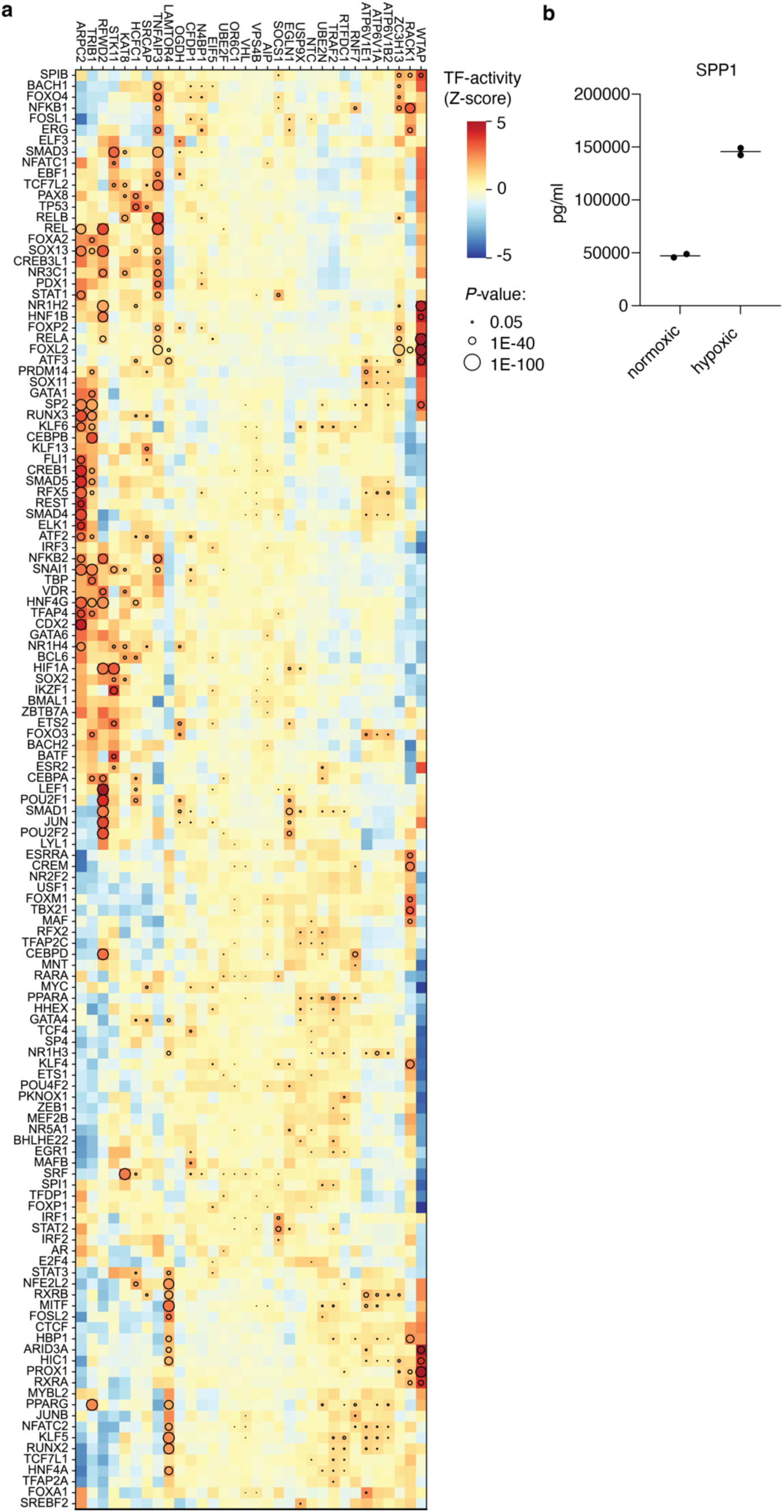
Regulon analysis on perturbed hMDMs. **a,** TF activity scores (color bar) of TFs (rows) in perturbed hMDMs (column) upon IFN-γ priming **b**, hMDMs cultivated in normoxic or hypoxic conditions for 24h, followed by SPP1 measurements in supernatants using Luminex. Each symbol represents an individual measurement of one donor (n=2).

**Supplementary Fig 6:**
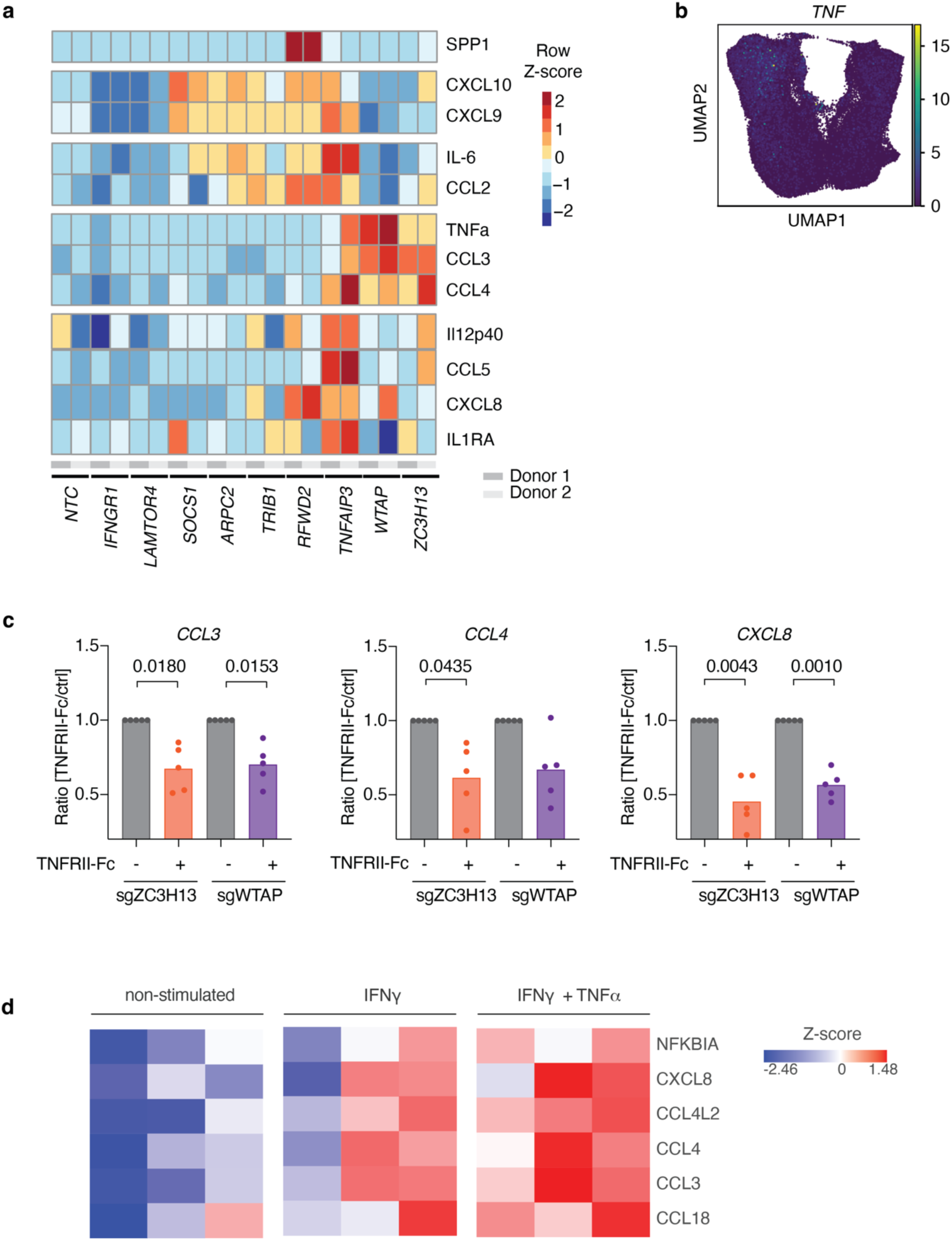
Additional cytokine secretion analysis. **a,** Heatmap as shown in Fig. 3a with additional cytokine and chemokine values upon IFN-γ stimulation. Z-score scaling for each cytokine or chemokine shown for two individual donors. **b**, *TNF* expression in UMAP projection. **c**, Gene expression of *CCL3*, *CCL4*, and *CXCL8* assessed by qPCR of perturbed hMDMs 18 h after IFN-γ priming with and without addition of TNFRII-Fc in culture medium; each symbol represents an individual donor (n=5). Summary data of 3 individual experiments are shown as mean, with *P* values determined by ANOVA with Sidak’s multiple comparisons test. **d**, Heatmap showing P03 genes in non-stimulated, IFN-γ primed and IFN-γ and TNFα co-stimulated cells in hMDMs, each column represents an individual donor (n=3).

**Supplementary Fig 7:**
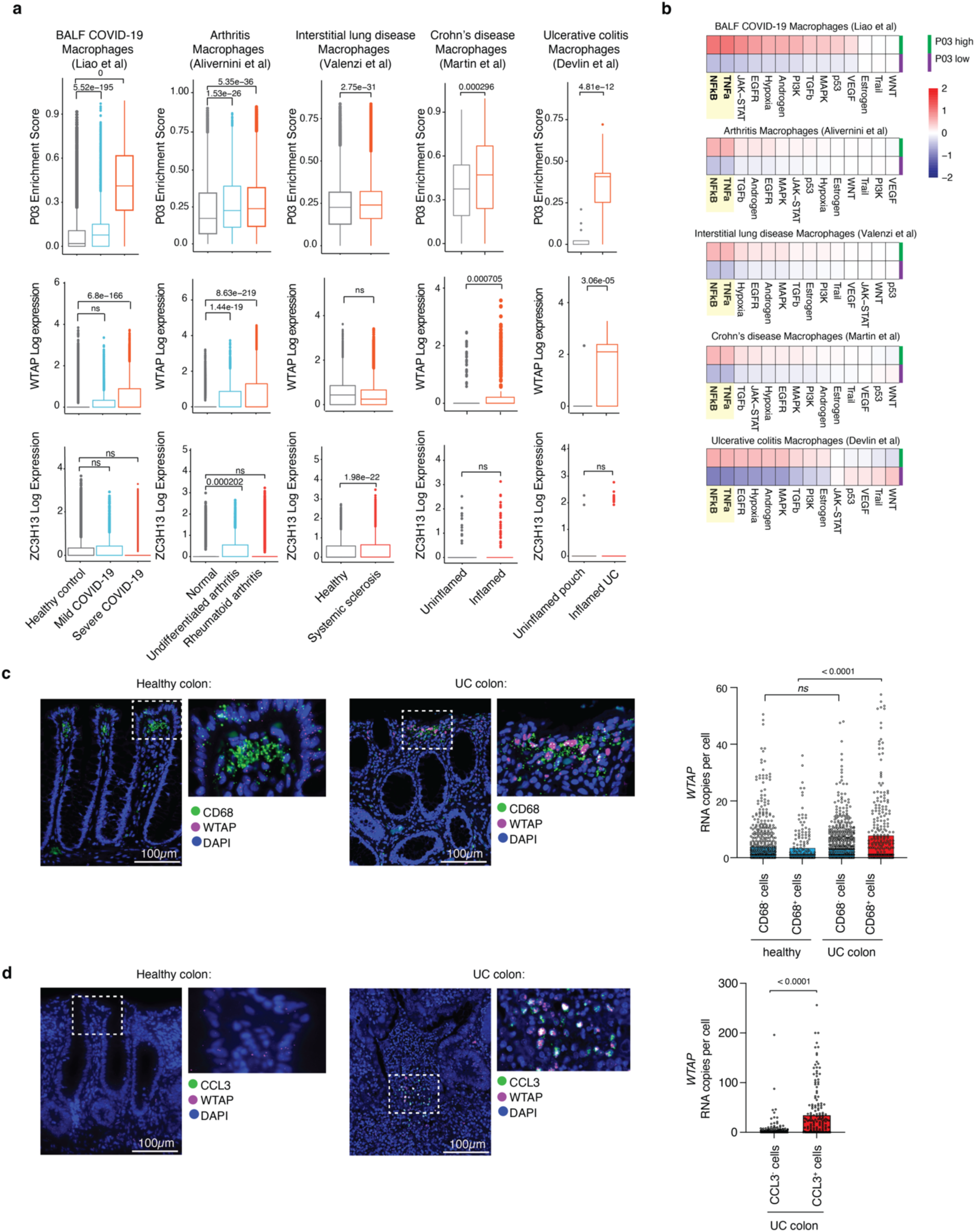
WTAP associates with P03, a TNFα-driven inflammatory macrophage state across disease conditions. **a,** P03 enrichment scores, WTAP and ZC3H13 expression in the macrophage compartment are shown in healthy BALF (n=3), mild (n=3) and severe (n=5) COVID-19^61^, normal synovium (n=4) undifferentiated arthritis (n=3) and arthritis^62^, (n=19), healthy (n=6) and systemic sclerosis (n=10)^63^, uninflamed (n=10) and inflamed CD (n=11)^64^ as well as uninflamed pouch (n=3) and inflamed UC (n=10)^65^. Box plots summarize the median, interquartile and 75% quartile range for each tissue type. *P* values determined by two-sided t-test. **b,** Pathway enrichments scores for P03-high vs. -low macrophages for disease indication as described in (a, upper row). High vs low P03 groups were defined using the median enrichment score in macrophages for each study. **c,** Representative RNA *in situ* hybridization of *WTAP* and *CD68*, a known macrophage marker (**c**) or *CCL3*, a gene in P03 (**d**) in healthy and ulcerative colitis colon tissues. Images on the right are higher magnification of images on the left. (right) Quantification of *WTAP* expression in *CD68* (**c**) or *CCL3* (**d**) expressing and non-expressing cells. Each dot represents the gene expression value from one cell (n=3 healthy donors) (n=4 UC patients). Scale bar = 100 μM. Summary data are shown as mean, with *P* values determined by Tukey’s multiple comparison (**c**) and Student two-sided t-test (**d**).

**Supplementary Fig 8:**
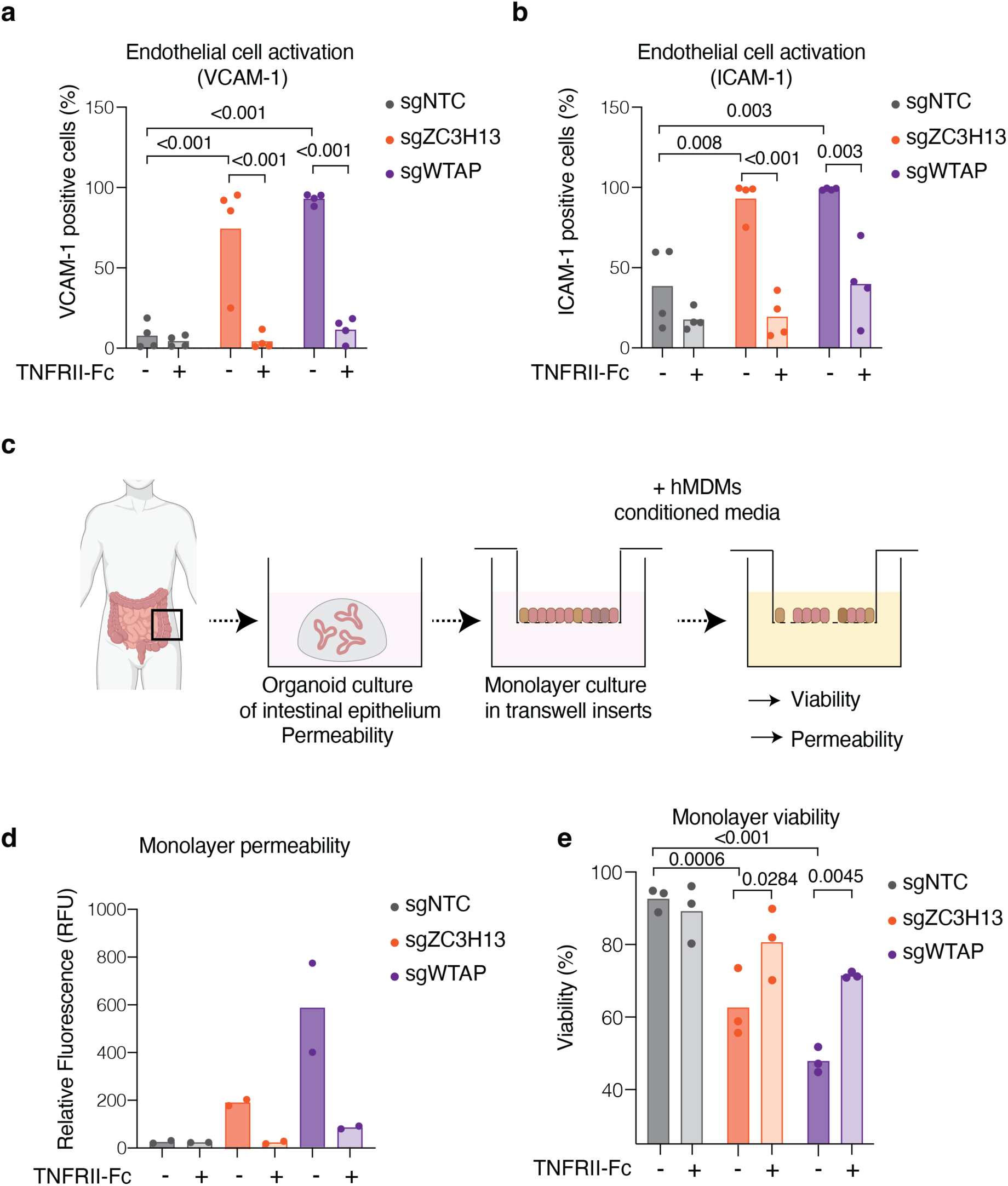
*ZC3H13-* or *WTAP*-perturbed hMDMs promote TNFα-dependent paracrine effects. **a,** Flow cytometry quantification of VCAM-1 and **b**, ICAM-1 positive endothelial cells upon treatment with conditioned macrophage media in presence or absence of TNFRII-Fc as indicated. **c,** Experimental workflow depicting intestinal barrier integrity measurements **d,** barrier permeability (FITC-dextran leakage across monolayer) upon treatment with conditioned macrophage media and **e**, Viability of treated monolayers assessed (MTA assay). a, b, d, e, Each symbol represents an individual donor (a and b, n=4) (d, n=2) (e, n=3). Summary data are shown as mean, with *P* values determined by ANOVA with Sidak’s (a, b, e) multiple comparisons test.

**Supplementary Fig 9:**
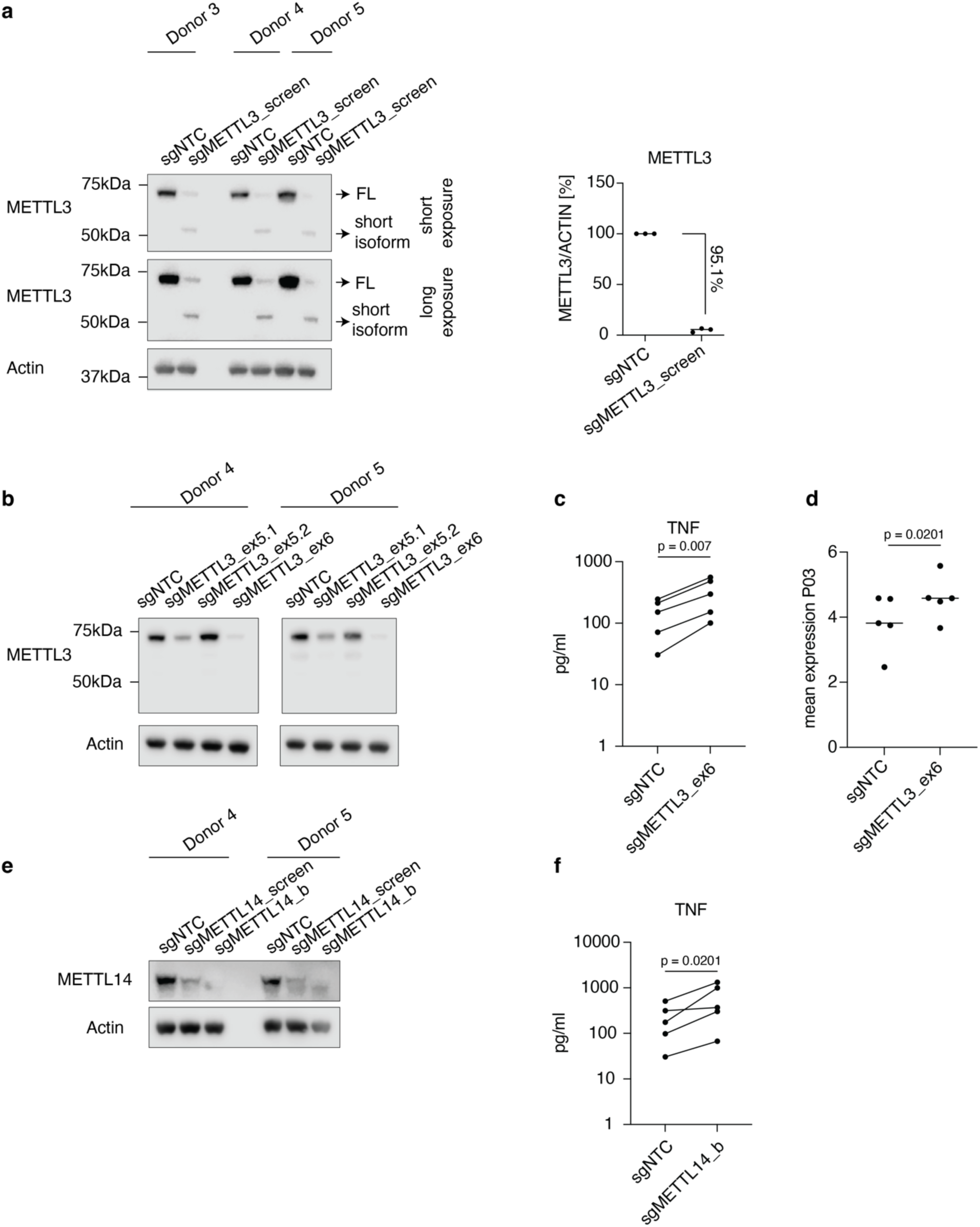
Perturbation of METTL3 and METTL14 in hMDMs elevate TNFα secretion. **a,** Western blot of METTL3 perturbed with sgRNAs utilized in scCRISPR screen (sgMETTL3_screen) and quantification of full length METTL3 (FL) (right). **b,** Western blot of METTL3 perturbed with sgRNAs targeting exon 5 (sgMETTL3_ex5.1 and sgMETTL3_ex5.2) and exon 6 (sgMETTL3_ex6). **c,** TNFα cytokine release and **d**, mean P03 of METTL3 perturbed cells using sgMETTL3_ex6. **e**, Western blot of METTL14 perturbed with sgRNAs utilized in scCRISPR screen (sgMETTL14_screen) and (sgMETTL14_b). **f,** TNFα cytokine release in METTL14 perturbed hMDMs using sgMETTL14_b. a, c, d, f, Each symbol represents an individual donor (a n=3) (c, d, f n=5). Summary data are shown as mean, with *P* values determined by and Student two-sided t-test (c, d, f). a, representative Western blot image of 3 donors, b and e of 2 donors.

**Supplementary Fig 10:**
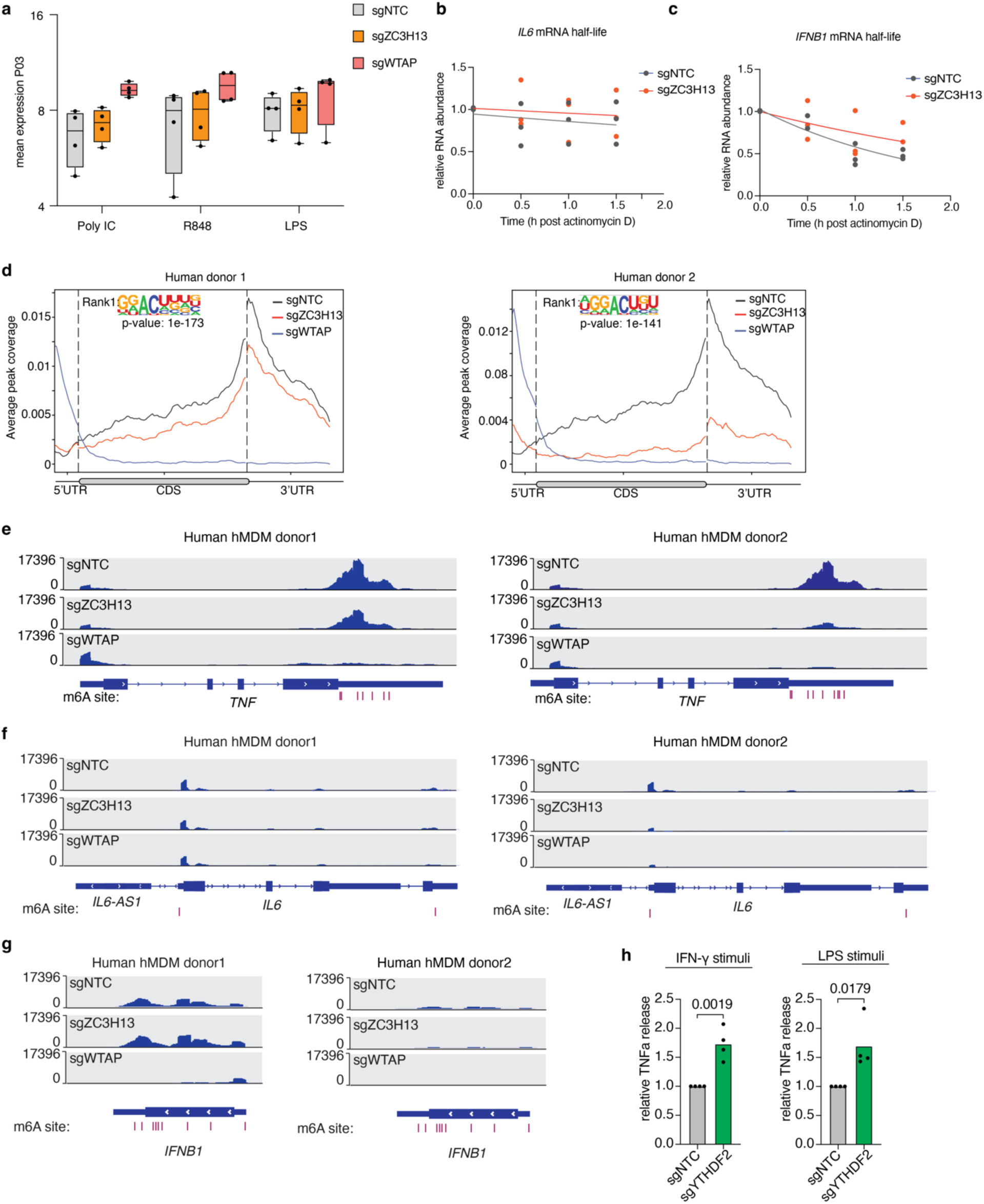
Genomic visualization of m6A-eCLIP. **a,** Mean expression of P03 in NTC-, *WTAP*- and *ZC3H13*-perturbed hMDMs across indicated treatment conditions. **b, c,** Relative *IL6 and IFNB1* mRNA levels upon ActinomycinD treatment in *ZC3H13*-depleted and NTC control cells. **d**, Normalized distribution of m6A peaks across 5’ UTR, CDS, and 3’UTR of perturbed hMDMs as indicated for two individual donors (1 and 2). Top motif enriched in m6A-eCLIP peaks derived from NTC control cells. **e**, Genomic visualization of normalized m6A-eCLIP signal along *TNF*, *IL6* (**f**) and *IFNB1* (**g**) in perturbed hMDMs as indicated for two individual donors (1 and 2); vertical red lines depict m6A residues **h**, TNFα cytokine levels determined by Luminex analysis of cell culture media of perturbed hMDMs as indicated 18 h after LPS or IFN-γ stimulation. a, b, c and h Each symbol represents an individual donor (a and h, n=4) (b and c n=3). h, *P* values determined by Students two-sided t-test.

**Supplementary Fig 11:**
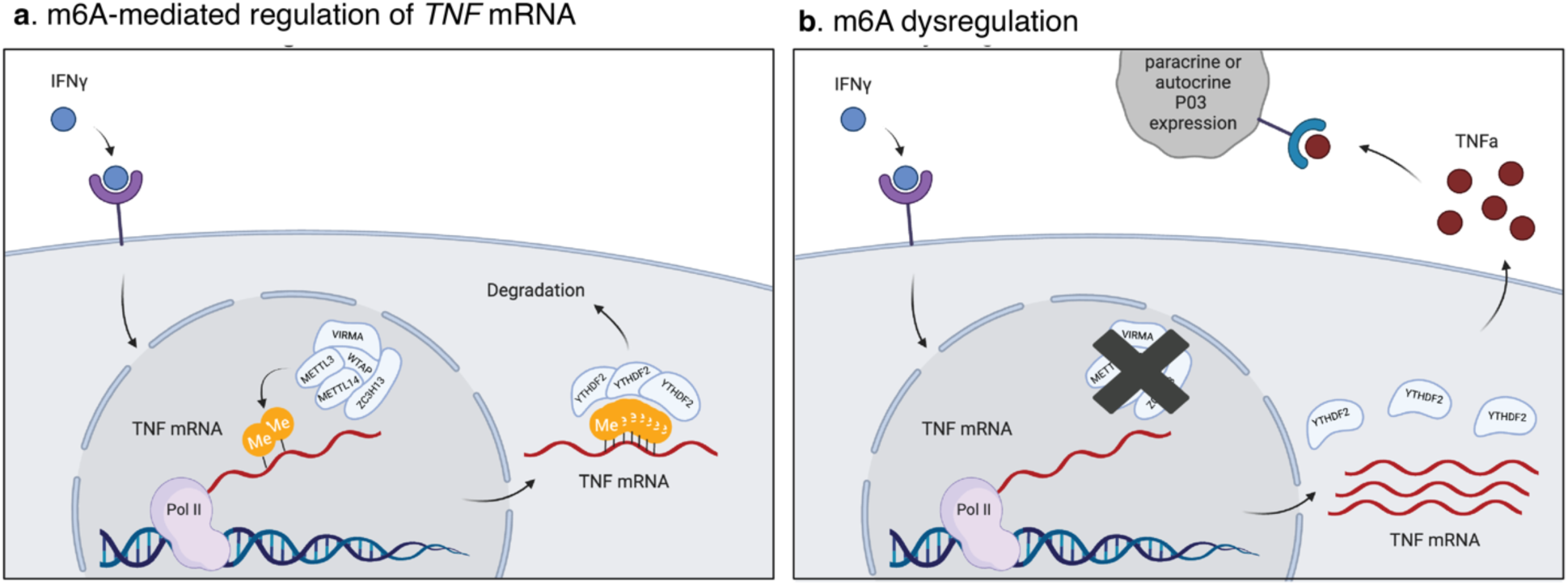
m6A mediated regulation of *TNF*. **a**, IFN-γ priming induces low level expression of *TNF* transcript, which becomes m6A modified by the m6A writer complex. Installed m6A can be detected by m6A reader proteins such as YTHDF2, which promote degradation of the transcript. **b**, Depletion of individual m6A writer components leads to loss of m6A installation. Unmodified *TNF* mRNA cannot be recognized by m6A reader proteins, resulting in accumulation of TNFα transcript and consequently, increased protein expression. Elevated levels of TNFα induce P03 expression likely through paracrine and/or autocrine signaling via TNF receptors.

## ACKNOWLEDGEMENTS

We are grateful for the cooperation of Donor Network West and all of the organ and tissue donors and their families, for giving the gift of life and the gift of knowledge, by their generous donation. We thank members of the Next Generation sequencing Facility, Isabelle Lehoux and Zhong Rong Li for support in construct design; Eclipse BioInnovations Inc. for performing m6A-seq and analysis; Lucia Taraborrelli, Ricardo Mendes Leitao, Junghyun Lim and members of the Cancer Immunology and Immunology Discovery departments at Genentech for technical and intellectual support.

## AUTHOR CONTRIBUTIONS

S.M.H., S.X., C.E., J.K., J.L, M.C., L.H., and C.C designed and performed experiments, analyzed and interpreted data. S.M.C, R.N.P and S.Z.W. analyzed and interpreted data. A.L, J.P.F, M.C. analyzed genome-wide CRISPR-screen data sets. B.Y. and H. H. performed PheWAS analysis. E.F. and A.N provided reagents. S.M., M.K. K.G.S., provided essential conceptual input. S.M.H and A.M. conceptualized the study. S.M.H wrote the manuscript with input from all authors. S.J.T. and A.M. oversaw the project.

## METHODS

### Mice

C57BL/6N animals were obtained from Jackson labs. Protocols were approved by the Genentech Institutional Animal Care and Use Committee. Studies were executed by following relevant ethical regulations as detailed in associated animal use protocols.

### mBMDM culture

mBMDM were cultured in DMEM high glucose supplemented with 10% FBS (F2442, Sigma), 2mM Glutamax (35050061, Thermofisher), 100 U/ml Penicillin-Streptomycin (Thermofisher, 15140122), 50 ng/ml M-CSF (Genentech).

### Cell stimulation

Cells were stimulated with ultrapure LPS (5 ng/ml unless specified otherwise) (E.coli 0111:B4, Invitrogen), recombinant human IFN-γ (2 ng/ml) (285-IF-100, R&D Systems), recombinant human TNFα (10 ng/ml) (10291-TA-020, R&D Systems), poly I:C (1µg / ml) (tlrI-pic, Invitrogen), R848 (1µg / ml) (tlrI-r848, Invitrogen), recombinant mouse IFN-γ (20 ng/ml) (485-MI-100, R&D Systems), recombinant mouse TNFα (5 ng/ml). (410-MT-010/CF, R&D Systems), recombinant mouse IL1b (20 ng/ml) (401-ML-005/CF, R&D Systems) at indicated time points and concentrations. In vitro TNFα-blocking studies were performed using recombinant human TNFRII/TNFRSF1B-Fc chimera protein (1 µg / ml) (726-R2-050, R&D Systems) for 18 h.

### Generation of lentiviral particles

Lentiviral particles were produced and titered as previously described^66^. Infectious virus titer was obtained by culturing and infecting BMDM plated in 12 well plates (Falcon, 351143) following a scaled version of the culture and infection protocol for the large-scale lentivirus library infection. Infectious particles/ml were extrapolated using virus dilutions and measuring resistance to puromycin (A11138-03, Thermofisher) at a concentration of 5 µg/ ml for each lentivirus library pool.

### Pooled genome-wide CRISPR screen in mBMDMs

On Day 1, an estimated 682 million cells isolated from the femoral and tibial bone marrow of 75 13-week-old mice were distributed to 80 T-175 flasks (Corning, 431466) containing 20 ml of BMDM culture medium. After 48 h (or day 3), unattached cells were washed away with PBS allowing ∼1.2 billion cells to be staged as 400 million cells in three replicates, 26 flasks each for infection with 8 combined lentivirus sgRNA library pools expressing 152 339 sgRNA targeting 18776 genes, control genes and non-targeting sgRNA at an MOI of 0.5 to achieve a representation of at least 1500 cells per guide. Transduced BMDMs were harvested on day 6 by scraping the culture flasks in PBS, setting aside ∼120 million cells of each replicate for reference genomes while the remaining cells, ∼60 million for each replicate were electroporated with Cas9-ribonuclear particles (Cas9-RNP) as previously described^67^. Briefly, non-targeting control (NTC) sgRNA comprised of gNTC Alt-R® CRISPR-Cas9 Negative Control crRNA #1(1072544, Integrated DNA Technologies) and Alt-R tracrRNA (Alt-R® CRISPR-Cas9 tracrRNA (1072534, Integrated DNA Technologies) used as a swap guide was complexed with Alt-R® S.p. HiFi Cas9 Nuclease V3 (1081061, IDT) at a 3:1 molar ratio at room temperature for 20 min^68^. Harvested BMDMs were resuspended in P3 primary cell nucleofection buffer (V4XP-3024, Lonza) and electroporated with generated Cas9-RNP complexes using supplied nucleofector cuvettes (Lonza) in a 4D-Nucleofector X Unit. Cells were then replated for continuous culture. On day 7 puromycin was introduced to the culture medium. Thereafter, one half of the culture media was replaced daily for fresh culture medium by puromycin (5 µg/ml) supplemented media for 7 consecutive days. On day 14, cells were stimulated with 20 ng/ml mIFN-γ for 18 h in BMDM media without rmM-CSF. The next day ∼ 700 million stimulated BMDMs were harvested for each replicate, washed, fixed and stained at a density of 10 million cells/100ul. Staining proceeded with an initial Fc-block step (BD553142, BD bioscience) combined with fixable viability dye eFluor 780 (65-0865-14, eBioscience). cells were stained with anti-CD45 BV510 (BD563851, BD bioscience), anti-CD11b BUV737 (BD564443, BD bioscience), anti-F4/80 PerCpCy5.5 (123127, BioLegend), followed by fixation and intracellular staining with anti-iNOS FITC (53-5920-82, eBioscience). Staining steps proceeded with incubation, centrifugation and washing at each step. ∼ 100 million fixed cells of each replicate were kept as an unsorted reference. Cells were sorted by gating on live, CD11b +, F4/80 +, CD45 + and gated on the top 10% iNOS expressing cells using a BD FAC Symphony S6 Cell Sorter resulting in ∼ 6 million cells for each bin and replicate.

### Library generation for sgRNA quantification of genome-wide CRISPR screen

Genomic DNA was extracted using Gentra Puregene Cell Kit (158388, Qiagen) using extended incubation at 55 °C with Proteinase K (19131, Qiagen) supplemented in the lysis buffer (25 µl per 10 ml) to reduce any DNA-protein cross-links. DNA was allowed to dissolve for 48 h prior to quantification using Qubit dsDNA Quantitation range kit (Q32853, Thermofisher). From the estimated 3 to 6 million cells in each of these bins, 20 to 40 % of genomic DNA was recovered by mass. sgRNA guide sequences were recovered as amplicons generated by PCR of the genomic DNA using Phusion High-Fidelity PCR Master Mix with GC Buffer (M0531L, NEB) with forward (TCTTGTGGAAAGGACGAGGTACCG) and reverse (TCTACTATTCTTTCCCCTGCACTGT) primers representing complimentary guide flanking sequences. Adapter ligation and indexing PCR was performed on these amplicons using KAPA HTP Library Preparation Kit (KK8234, Kapa biosystems) for multiplexing and sequencing on an Illumina HiSeq5 platform.

### Bioinformatics and statistics methods of pooled-genome wide CRISPR screen in mBMDMs

Amplicons were sequenced on an Illumina HiSeq 4000 instrument, using 50 bp paired-end reads. The average Phred quality score was above 38 for all 18 samples, and all 18 samples had at least 150 M reads. In each read pair, both read sequences were searched for an exact match of any one of the gRNA barcode sequences, using an in-house software package written in R/C++. The number of reads matched to each guide in each sample was used to obtain a guide-by-sample count matrix for further analysis.

The raw count data were stored in a standard Bioconductor summarized experiment object^69^. Normalization was performed to adjust for the difference in sequencing depth between samples, and to account for potential compositional biases due to the competitive nature of pooled genetic screening. We estimated normalization factors for each sample by applying the TMM method^70^ on the count data using gRNAs targeting a gold-standard set of non-essential genes (obtained from^71^). We performed a differential abundance analysis at the gRNA level using the popular limma-voom approach^72^. Specifically, we fitted a linear model to the log-CPM values for each gRNA, using voom-derived observation and quality weights. We performed robust empirical Bayes shrinkage to obtain shrunken variance estimates for each gRNA, and we used moderated F-tests to compute *P* values for each of the two-group comparisons of interest. To control the FDR in each comparison, we applied the standard Benjamini-Hochberg method to obtain an adjusted *P* value for each gRNA. We then used the gRNA-level statistics to obtain gene-level summaries using two complementary statistical approaches. The first approach was to aggregate gRNA statistics by reusing the “fry” gene-set enrichment analysis method implemented in limma, and considering gRNAs targeting a given gene as a “gene set”. This allows the detection of genes that are consistently enriched or depleted for the majority of the gRNAs designed for each gene. The second approach uses Simes’ method to obtain a combined *P* value for each gene based on the *P* values for all associated gRNAs^73^. This allows the detection of differentially-abundant genes in cases where only a small proportion of the gRNAs show a strong signal. Gene-level *P* values were corrected for multiple comparisons using the Benjamini-Hochberg method. For both methods, hits were obtained using a FDR threshold of 5%.

### Endothelial cell culture

HUVECs were obtained from Lonza and cultured in EGM-2 Endothelial cell growth medium-2 bullet kit (Lonza, CC-3162) supplemented with penicillin-streptomycin and glutamine on gelatin coated plates (Millipore Sigma, ES-006-B). For activation endothelial cells were incubated with hMDM conditioned media for 6 h in absence or presence of TNFRII/TNFRSF1B-Fc chimera protein (726-R2-050, R&D Systems) at 1 µg/ml.

### Intestinal crypt isolation and organoid culture

Crypt isolation and organoid culture was performed as previously described^74^ with minor modifications. Briefly, large intestine tissue from normal donors was acquired from Donor Network West. Derived mucosal layer was washed, divided into 1x1 inch pieces and epithelial lavers were separated from the muscle using forceps. Obtained epithelium was wash in (DMEM high glucose containing with 1x Penicillin-Streptomycin (Gibco, 15140-122), 10mM HEPES (Gibco, 15630130), 1x GlutaMAX Supplement (Gibco, 35050061) and 2.5 ug/mL Amphotericin B (Gibco, 15290018)) and cut into tiny pieces (∼2 mm^2^) with scalpels. To release crypts the washed epithelium was treated with 20 mL of 2.5 mM EDTA in PBS containing with 10 uM Y27632 (Stemcell, 72304) at 37°C in an end over end rotator for 12 min. Derived crypts were filtered through a 100 μM filter and rigorously washed. The resulting clean crypts were suspended in Matrigel (Corning, 354234) and grown in droplets overlayed with intestinal growth media containing Advanced DMEM/F12 (Gibco, 12634010), 1x Glutamax (Gibco, 35050061), 1x Penicillin-Streptomycin (Gibco, 15140122), 10 mM HEPES (Gibco, 15630130), 10 mM Nicotinamide (Sigma Aldrich, N0636), 1x B27 (Thermofisher Scientific, 12587-010), 1x N2 (Thermofisher, 17502048), 100 ug/mL Primocin (InvivoGen, ant-pm-1), 10 µM Y27632 (Stemcell Technologies, 72304), 100 ng/mL IGF-1 (Biolegend, 590904), 50 ng/mL FGF2 (Peprotech, AF-100-18B), 50ng/mL EGF (R&D Systems, 236-RG), 100 ng/mL Noggin (R&D Systems, 6057-NG), 500 nM A83-01 (Tocris, 2939), 10 nM Gastrin (Tocris, 3006), 250 ng/ml R-spondin 3 (Genentech) and 0.2 nM Wnt Surrogate (ImmunoPrecise, N100).

To establish monolayers, grown organoids were harvested and treated with TrypLE (Gibco, A1217701) to obtain single cells. 0.3 million cells were plated on a collagen (Corning, 354236) coated transwell (Costar, 38024) and grown for 5 days in intestinal growth media or until confluent. Derived monolayers were subsequently differentiated for 4 additional days in intestinal growth media to which 10 µM DAPT (Sigma, D5942) and 1 µM PD0325901 (Sigma, PZ0162) was added and Wnt Surrogate adjusted to 0.02 nM. To assess cytokine mediated effects of perturbed macrophages on intestinal epithelial cells, differentiated monolayers were treated for 3 days with conditioned macrophage media. Conditioned media was derived from IFN-γ (0.2 ng/ml) stimulated NTC-, *ZC3H13*- and *WTAP*-perturbed hMDMs 18 h after stimulation.

On day 3 of treatment, monolayer permeability was assessed by transferring the transwell plates in a new receiver plate containing 200 µl Hank’s balanced salt sodium (HBSS) (Gibco 14025-092). To determine permeability of the monolayers (1 mg/ml) FITC-dextran (Sigma-Aldrich FD4-100MG) in HBSS was added to the apical side for 2 h. FITC-dextran was measured in the receiver plated at ex:494 and em:521.

On day 3 of treatment, cell viability of intestinal monolayers was determined using MTS assay kit (ab197010). MTS reagent was added to cell culture media to the transwell plate as described by the manufacturer in standard culture conditions for 4 h. Absorbance was determined at 490 nm.

### Donors for PBMC isolation

Human blood samples were obtained from healthy volunteers through the Genentech employee donation program, study protocols were reviewed and approved by the Western Institutional Review Board.

### Human MDM culture

PBMCs were isolated from the blood of healthy volunteers using Lymphoprep medium (STEMCELL Technologies). Miltenyi human classical monocyte cell isolation kit (130-117-337) and Miltenyi LS column (130-042-401) were used for monocyte isolation. Macrophage differentiation was induced by culturing monocytes at a density of 1 x 10^6^ cells/ml in DMEM high glucose plus 10 % FBS plus penicillin-streptomycin plus 50 ng/ml rhGM-CSF for 9 days. Adherent macrophages were stimulated as indicated.

### Bulk RNA-seq

Total RNA was isolated using the Qiagen RNeasy isolation kit (74104) and quantified with Qubit RNA HS Assay Kit (Thermo Fisher Scientific) and quality was assessed using RNA ScreenTape on 4200 TapeStation (Agilent Technologies). For sequencing library generation, the Truseq Stranded mRNA kit (Illumina) was used with an input of 100 nanograms of total RNA. Libraries were quantified with Qubit dsDNA HS Assay Kit (Thermo Fisher Scientific) and the average library size was determined using D1000 ScreenTape on 4200 TapeStation (Agilent Technologies). Libraries were pooled and sequenced on NovaSeq 6000 (Illumina) to generate 30 millions single-end 50-base pair reads for each sample.

RNA-seq data were analyzed using HTSeqGenie (https://bioconductor.org/packages/release/bioc/html/HTSeqGenie.html) as follows: first, reads with low nucleotide qualities (70% of bases with quality <23) or matches to ribosomal RNA and adapter sequences were removed. The remaining reads were aligned to the human reference genome (hg38) using GSNAP, allowing a maximum of 2 mismatches per 75-base sequence. Subsequently, the number of fragments was calculated by counting the number of uniquely mapped concordant pairs to the exons of each RefSeq gene using the *summarize Overlaps* function from the GenomicAlignments package^75^. Expression levels of each gene were normalized to Reads Per Kilobase of transcript, per Million mapped reads (RPKM) and log2 transformed. Mean expression values of P03 (*NFKBIA, CXCL8, CCL4L2, CCL4, CCL3, CCL18*) were determined using Partek Flow.

### Ethics

Written informed consent was obtained in accordance with research and ethics committee (REC) (Newcastle and North Tyneside 1 REC 10/H0906/41 and Newcastle and North Tyneside 2 REC 22/02; Mayo Clinic Institutional Review Board 10-006628) approval. These studies were performed according to the principles of the Declaration of Helsinki. Peripheral blood samples from healthy male and female donors were provided by the Samples4Science donor program at Genentech. Donors provided written informed consent and sample collection was approved by the Western Institutional Review Board.

### CRISPR Cas9-mediated deletion in human and mouse macrophages

sgRNAs sequences were selected using either the IDT pre-design guide tool (https://www.idtdna.com) or an in-house algorithm. Chosen sequences were obtained as Alt-R® CRISPR-Cas9 sgRNA from IDT and reconstituted to 100 µM with nuclease-Free Duplex Buffer (11-01-03-01, IDT). For efficient deletion of target genes in hMDMs, 4 sgRNAs targeting the same gene were mixed in equimolar ratios in a sterile PCR strip, unless indicated otherwise. For follow up studies in BMDMs 2 sgRNAs targeting the same genes were combined. As negative controls, non-targeting sgRNAs (NTC), as well as sgRNAs against the “safe harbor locus” AAVS1 and the non-essential gene OR6C1 were chosen (https://depmap.org/portal/gene/OR6C1?tab=overview).

For generating of Cas9-RNPs, sgRNAs were complexed with Alt-R® S.p. HiFi Cas9 Nuclease V3 (1081061, IDT) at a 3:1 molar ratio (3 µl sgRNAs + 1 µl Cas9 Nuclease) at room temperature for 20 min for each reaction. Electroporation was performed as described in Freund et al^23^. Briefly, 3 x 10^6^ freshly isolated monocytes or BMDMs were resuspended in 20 µl of P3 primary cell nucleofection buffer (Lonza). Cells were then added to the Cas9-RNP complexes, mixed carefully and immediately loaded into the supplied nucleofector cassette strip (Lonza). Electroporation was performed using a 4D-Nucleofector X Unit, program: CM-137. After the reaction cells were carefully harvested with prewarmed media and cultured as described above.

### Generation, processing and analysis of scCRISPR screen data

Perturbed human monocyte-derived macrophages were generated and differentiated as described above in an arrayed fashion. On day 9 cells were stimulated with 0.2 ng/ml hrIFN-γ or fresh medium as control for 18 h. Cells were harvested using Detachin (Genlantis), washed in PBS and resuspended in cell-staining buffer (BioLegend) containing FcX blocking antibody (Biolegend). To achieve perturbation specific labeling of cells, each knock-out was stained with a unique TotalSeq^TM^-A hashtag oligo (HTO) 1-10 (BioLegend) combination, pooled and encapsulated using a Chromium Single Cell Controller (10X Genomics). Each lane was loaded with 20 000 cells and libraries were prepared following the Chromium Single Cell 3′ Library and Gel Bead Kit v2, following the manufacturer’s manual (CG00052 Chromium Single Cell 3′ Reagent Kits v2 User Guide RevA; 10X Genomics). HTO library was prepared according to published guidelines^76^. Libraries were profiled by a Bioanalyzer High Sensitivity DNA kit (Agilent Technologies) and quantified using the Kapa Library Quantification Kit (Kapa Biosystems). Each library was sequenced in one lane of HiSeq4000 (Illumina) following the manufacturer’s sequencing specification (10X Genomics).

For processing of 10X gene expression and HTO libraries, cell by gene, and cell by feature barcode count matrices were generated using the CellRanger workflow from cumulus (version 0.14.0). Gene expression reads were aligned against the GRCh38 reference genome, and feature barcodes against TotalSeq^TM^-A sequences. Feature barcode demultiplexing was performed using the demultiplexing workflow from cumulus (version 0.14.0) which uses DemuxEM. Cells identified as doublets by DemuxEM were excluded from analysis. We further removed cells with less than 500 detected genes, as well as cells where more than 20 % transcript counts derived from mitochondrial-encoded genes. Because high UMI counts might indicate doublets, we excluded cells with more than 6000 UMI counts.

The computational analysis for the filtered and quality-controlled cells was largely performed using the python package Scanpy using standard protocols. Briefly, count matrices were merged, normalized and logarithmized before selecting the top 2000 highly variable genes per donor with a mean expression between 0.01 and 5. Harmony was used to integrate the datasets by donor, and this new projection used to calculate the neighborhood graph and UMAP embedding. We sequenced a total of 248,262 raw cells, and 176,564 high quality cells with a median of 4,537 cells per genetic perturbation were retained for further analysis after standard quality control. Perturbation of *RBBP6* and *RAN* in hMDMs led to a drastic reduction in cell numbers in both donors and were therefore excluded from further analysis.

Topic analysis was performed on raw counts using the Latent Dirichlet Allocation (LDA) implementation from Scikit Learn using 17 components, a prior of document topic and topic word distribution of 0.1 and 0.001, respectively. To detect enrichment of perturbations in each topic, the cohen-d effect size was calculated per topic between the NTC and each perturbation using the statistical python library Pingouin. Regulon activity was calculated using the python implementation of DoRothEA using the non-academic version of its regulons^77,78^. TF activities were calculated by fitting a multivariate linear model, and then ranked using a Wilcoxon test. Energy distance test for identifying perturbations that induced altered transcriptional states was performed as previously described^79^ using 1000 permutations.

Published datasets were processed in a similar way as our dataset. We scored P03 (*CCL3*, *CCL4*, *CCL18*, *CCL3L1*, *CCL4L2*, *CXCL8*, *NFKBIA*) macrophages extracted from independent scRNA-Seq datasets^61–65^. Raw or normalized count matrices were pulled from each respective study and processed using Seurat v4.1.1 using default parameters recommended by the developers. Gene signatures were then scored in the transformed expression matrix of single cells using AUCell v1.19.1^80^. Statistical significance between groups was determined using a two-sided t-test in a comparison of mean Area Under Curve (AUC) values between groups. High vs low P03 groups were defined using the median enrichment score in macrophages of each study. Pathway activity for published scRNAseq datasets was determined using PROGENy^81^.

### Cytokine measurements

Cytokines and chemokines from culture supernatants were analyzed using human cytokine MILLIPLEX Luminex (EMD Millipore). Briefly, 25 μl of sample and 25 μl of analyte-specific color-coded magnetic beads coated with capture antibodies were mixed. Biotinylated detection antibodies were added followed by incubating with streptavidin-phycoerythrin. The median fluorescence intensity (MFI) data were analyzed using a five-parameter logistic method for calculating cytokine/chemokine concentrations in samples.

### Flow Cytometry

BMDMs were incubated in Fc block (BD biosciences, 24G2) and fixable viability dye eFluor 780 (Invitrogen) for 15 min in PBS at 4 °C. Cells were washed once and stained with the following antibodies: anti-CD11b (BD Horizon, M1/70), anti-iNOS (Invitrogen, CXNFT) and anti-TNFα (Invitrogen, MP6-XT22).

Endothelial cells (HUVEC) were incubated in Fc block (BD biosciences, 564219) and fixable viability dye eFluor 780 (Invitrogen) for 15 min in PBS at 4 °C. Cells were washed once and stained with the following antibodies: anti-VCAM-1 (BD biosciences 551146, clone 51-10C9) anti-ICAM-1 (BD biosciences 551146, clone HA58-10C9).

### RNA extraction and qPCR

A total of 1 x 10^6^ cells were lysed using RLT buffer (QIAGEN) and total RNA was extracted using RNeasy isolation kit plus (QIAGEN) according to manufacturer’s instructions. Obtained RNA was reverse-transcribed into cDNA using (iSCRIPT, BioRad) according to manufacturer’s instructions. qPCR was performed using TaqMan Gene Expression solutions using a QuantStudio Real-Time PCR System (Life Technologies). *RPL36* or *CASC3* were used as house-keeping genes to normalize critical threshold (CT) values. Primers are commercially available: CASC3 (Hs00201226_m1), RPL36 (Hs03006033_g1), TNF (Hs00174128_m1), IL6 (Hs00174131_m1), IFNB1 (Hs01077958_s1), CCL3 (Hs00234142_m1), CCL4 (Hs99999148_m1), CXCL8 (Hs00174103_m1).

### TNF**α** expression assay in HEK293T cells

HEK293T cells were seeded in 12-well plates and transfected with wild-type or mutant human *TNF* + 3’UTR constructs using Lipofectamine 2000. Expression of target genes was induced through addition of doxycycline (1 µg/ml) and BrefeldinA (5 µg/ml) into culture media 24 h after transfection for indicated time points.

### RNA decay assays

Human MDMs were stimulated with LPS (100 ng/ml) for 3 h. To inhibit transcription, the medium was renewed and supplemented with Actinomycin D (Sigma Aldrich) at a final concentration of 5 µM. Total RNA was extracted at indicated time points and used for RT-qPCR.

### ISH-RNAscope tissue stains

Full thickness punch biopsies were collected post-surgery from IBD and healthy control patients and placed in 4 % formaldehyde, and shipped to Genentech. After arrival, samples were transferred to 70 % ethanol and then processed in paraffin for embedding. Once embedded, 5 uM sections were cut and processed for RNAscope in situ hybridization reagents and protocols from Advanced Cell Diagnostics (RNAscope multiplex fluorescent detection reagents v2, 323110). Briefly, embedded sections were allowed to dry in a 60 °C oven for 1 h. Sections were rehydrated in two washes of xylene for 5 min each followed by two washes in 100 % ethanol, one wash in 95 % ethanol, and one wash in 90 % ethanol, all for 2 min each. After rehydration, the samples were incubated in hydrogen peroxide, boiled in antigen retrieval buffer, and then digested with proteinase for 15 min at 40 °C. After digestion, the slides were washed twice for 1 min with ISH wash buffer and then hybridized with probes for 2 h at 40 °C. After hybridization, amplification steps were completed according to Advanced Cell Diagnostics protocol. After the final amplification, slides were incubated with Opal detection reagents (OP-001003 and OP-001004, Akoya bioscience) before mounting. Whole slide images were acquired on a Zeiss Axioscan Z.1 (Jena, Germany) with 20x objective, Hamamatsu Flash camera (Hamamatsu, Japan) Excelitas Xylis light source (PA), and DAPI/CFP/TRITC/Cy5 filters from Semrock (NY).

### Immunoblotting

Cells were lysed using 2 x Laemmli (BioRad) supplemented with 10 mM DTT followed by denaturation at 95 C for 5 min. Cell lysates were separated by SDS-PAGE 4 - 12 % gradient Bis-tris gel (Novex) followed by protein transfer to a PVDF membrane (BioRad) and antibody incubation. For immunoblotting anti-TNFα (Abcam, ab183218), anti-METTL3 (Cell Signaling, 86132), anti-METTL14 (Cell Signaling, 48699), anti-Actin (Cell Signaling, 5125) and goat anti-rabbit IgG-HRP (Cell Signaling Technology 7074, polyclonal) was used. ECL was recorded on an Azure Biosystems imager.

### PheWAS analysis

Phenome-wide association studies (PheWAS) enable the identification of phenotypes linked to individual genetic variants. Additionally, collapsing analysis, in which aggregation of rare variants is tested for association with disease, has emerged as a better way to understand the impact of rare variants captured in human disease. The UK Biobank Consortium updated the available Whole Genome Sequencing (WGS) data to a total of 490,640 individuals^56^ and made it available on the trusted research environment (Research Analysis Platform) on DNANexus in 2023. The SNPs and short Indels in the WGS dataset were jointly called over all individuals using GraphTyper^82^. We found 22 rare variants (see Table 1) in the WGS dataset (located in the interval: chr6:31577541-31577930) that were predicted to impact m6A installation on *TNF* mRNA. These SNPs were aggregated (collapsing analysis) into a single variable (i.e., carriers vs. non-carriers) which was then tested for association with the phenotypes. We grouped relevant ICD9/10 codes from UK Biobank into clinical meaningful phenotypes resulting in 1,695 phecodes (i.e., a group of ICD9/10 codes that capture clinical phenotypes). We then performed PheWAS for the 1,695 outcomes with the aggregated variable using Regenie. Regenie is a scalable and computationally efficient method for analyzing both binary and quantitative phenotypes^83^. It can handle imbalances in case-control ratios through firth correction. Regenie runs two steps: In Step 1, UKB array data was used as the input genetic data and a whole genome regression model is fitted on a predefined set of genetic markers. We applied the following quality control filters (as in ^83^) on the genotypic data to select variants that were kept in Step 1: 1) Minor allele frequency (MAF) ≥ %1, 2) Hardy-Weinberg Equilibrium (HWE) test < 1e-15, 3) genetic markers not present in low-complexity regions, 4) genotyping rate ≥ 99%, 5) genetic markers not involved in inter-chromosomal LD, and 6) LD pruning using a *R*^2^ threshold of 0.9 (window size = 1,000 ; step size = 100). In Step 2, we tested for association between the binary variable that represents the collapsed set of SNPs predicted to impact m6A installation on *TNF* mRNA and the phecode outcomes conditioned upon the prediction from the regression model in step 1. The following set of covariates were added to Step 1 and 2: *age*, *sex, age* × *sex, age*^2^, 20 ancestry principal components (PCs), and assessment center at which participants consented. The Regenie analysis was restricted to individuals with British ancestry (*n =* 424,136).

| Chr | Position (GRCh38) | REF | ALT | Allele frequency | Allele count | Number of samples | rsID | Presence in UKB imputed | Presence in UKB WES |
| --- | --- | --- | --- | --- | --- | --- | --- | --- | --- |
| 6 | 31577541 | G | A | 2.85E-05 | 28 | 490389 | rs190947828 | Yes | Yes* |
| 6 | 31577544 | C | T | 3.77E-05 | 37 | 490389 | rs752568182 | Yes | Yes* |
| 6 | 31577545 | G | A | 4.08E-06 | 4 | 490389 | rs758342189 | Yes | Yes* |
| 6 | 31577545 | G | C | 1.02E-06 | 1 | 490389 |  | No | Yes* |
| 6 | 31577548 | C | T | 0.000118273 | 116 | 490389 | rs777492465 | Yes | Yes* |
| 6 | 31577549 | A | G | 3.06E-06 | 3 | 490389 | rs1214395614 | No | Yes* |
| 6 | 31577549 | A | T | 1.02E-06 | 1 | 490389 |  | No | Yes* |
| 6 | 31577550 | T | A | 1.02E-06 | 1 | 490389 |  | No | Yes* |
| 6 | 31577672 | C | T | 1.02E-06 | 1 | 490366 | rs1398604432 | Yes | No |
| 6 | 31577704 | G | A | 1.02E-06 | 1 | 490356 |  | No | No |
| 6 | 31577706 | G | A | 1.02E-06 | 1 | 490357 |  | No | No |
| 6 | 31577708 | T | C | 1.02E-06 | 1 | 490356 | rs753952568 | No | No |
| 6 | 31577858 | G | A | 1.02E-06 | 1 | 490384 |  | No | No |
| 6 | 31577859 | G | A | 1.02E-06 | 1 | 490384 |  | No | No |
| 6 | 31577860 | A | G | 1.02E-06 | 1 | 490384 |  | Yes | No |
| 6 | 31577884 | T | C | 6.12E-06 | 6 | 490384 | rs892370128 | No | No |
| 6 | 31577887 | A | T | 1.02E-06 | 1 | 490386 |  | No | No |
| 6 | 31577888 | C | T | 1.02E-06 | 1 | 490387 |  | No | No |
| 6 | 31577894 | T | A | 1.02E-06 | 1 | 490386 | rs1262693118 | No | No |
| 6 | 31577896 | G | C | 1.02E-06 | 1 | 490387 | rs1238613915 | No | No |
| 6 | 31577926 | C | A | 1.02E-06 | 1 | 490319 |  | No | No |
| 6 | 31577930 | A | G | 1.02E-06 | 1 | 490320 |  | No | No |

\*These SNPs are part of the 100 bp flanking region of the targeted capture region. According to UK Biobank, only the targeted capture regions are required to meet all quality and coverage metrics.

### m6A-eCLIP and analysis

m6A-eCLIP was performed at Eclipse BioInnovations Inc, San Diego CA as described below. MDMs were stimulated with LPS (100 ng/ml) for 3 h. Total RNA was isolated from 20 x 10^6^ cells using Quick-RNA MiniPrep (Zymo) followed by a poly(A) selection using oligo (dT) beads (ThermoFisher) from 50 µg of total RNA. Obtained mRNA was DNase treated and immediately sheared into 100-200 nt fragments. After addition of an anti-m6A antibody (CST), samples were UV-C crosslinked using a UVP CL-1000 crosslinker at 2 rounds of 150 mJ/cm^2^ at 254 nm wavelength. The antibody-RNA complexes were then coupled overnight to protein G beads (CST). The next day, library preparation (including adapter ligations, SDS-PAGE electrophoresis and nitrocellulose membrane transfer, reverse transcription, and PCR amplification) was performed as previously described for standard eCLIP^84^, with the 30-110 kDa region size-selected by cutting from the membrane. 10 ng of fragmented mRNA was run as an RNA-seq input control, starting with FastAP treatment as described^84^. The final library’s shape and yield was assessed by Agilent Tape Station.

Due to the addition of 10 random nucleotides at the 5′ end serving as a unique molecular identifier (UMI)^85^ after sequencing primers are ligated to the 3′ end of cDNA molecules, UMIs were pruned from read sequences using umi_tools (v0.5.1)^86^. Next, 3′-adapters were trimmed from reads using cutadapt (v2.7)^87^, and reads shorter than 18 bp in length were removed. Reads were then mapped to a database of human repetitive elements and rRNA sequences compiled from Dfam^88^ and Genbank^89^. All non-repeat mapped reads were mapped to the human genome (hg38) using STAR (v2.6.0c)^90^. PCR duplicates were removed using umi_tools (v0.5.1) by utilizing UMI sequences from the read names and mapping positions. Peaks were identified within m6A-eCLIP samples using the peak caller CLIPper (https://github.com/YeoLab/clipper). For each peak, IP versus input fold enrichment values were calculated as a ratio of counts of reads overlapping the peak region in the IP and the input samples (read counts in each sample were normalized against the total number of reads in the sample after PCR duplicate removal). A *P* value was calculated for each peak by the Yates’ Chi-Square test, or Fisher Exact Test if the observed or expected read number was below 5. For single nucleotide calling 5′ mismatches were trimmed from read alignments and single nucleotide resolution crosslink sites were called from trimmed reads using the tool PureCLIP (v1.3.1)^91^. PureCLIP was run using the input control data in order to identify sites enriched in the IP over the input. Peaks and single nucleotide sites were annotated using transcript information from GENCODE (Release 35 or Release M25)^92^ with the following priority hierarchy to define the final annotation of overlapping features: protein coding transcript (CDS, UTRs, intron), followed by non-coding transcripts (exon, intron). Differential m6A-peaks between non-targeting control (NTC) and gRNAs were calculated using DESeq2. Metagene plots and de novo motif discovery were performed with the suite of tools for Motif Discovery and next-gen sequencing analysis HOMER (http://homer.ucsd.edu/homer/ngs/peakMotifs.html).

